# What works to improve species conservation state? An analysis of species whose state has improved and the actions responsible

**DOI:** 10.1101/2024.08.04.606320

**Authors:** Ashley T. Simkins, William J. Sutherland, Lynn V. Dicks, Craig Hilton-Taylor, Molly K. Grace, Stuart H.M. Butchart, Rebecca A. Senior, Silviu O. Petrovan

## Abstract

Understanding the consequences of past conservation efforts is essential to inform the means of maintaining and restoring species. Data from the IUCN Red List for 67,217 comprehensively assessed animal species were reviewed and analysed to determine (i) which conservation actions have been implemented for different species, (ii) which types of species have improved in state and (iii) which actions are likely to have driven the improvements. At least 51.8% (34,847) of assessed species have actions reported, mostly comprising protected areas (82.7%), with more actions reported for both terrestrial tetrapods and warm-water reef-building corals and fewer for fish, dragonflies and damselflies and crustaceans. Species at greater risk of extinction have a wider range of species-targeted actions reported compared to less threatened species, reflecting differences in documentation and conservation efforts. Six times more species have deteriorated rather than improved in their Red List category. Almost all species that improved have conservation actions in place; species that improved in state typically were historically at high risk of extinction, have smaller ranges and lacked a range of reported threats, particularly hunting and habitat loss or degradation. All types of conservation action were associated with improvements in state, especially reintroductions and invasive species control, alongside, for amphibians and birds, area management. This suggests a range of conservation interventions have successfully conserved some species at greatest risk but have rarely recovered populations to resilient levels. Scaling up the extent and intensity of conservation interventions, particularly landscape-scale actions that benefit broadly distributed species, is urgently needed to assist the recovery of biodiversity.

## Introduction

Humanity is inextricably dependent on the rest of biodiversity for the range of functions and ecosystem services it provides. However, we are facing a global biodiversity crisis, with 28% of 150,388 assessed species threatened with extinction (IUCN, 2023), and an estimated one million species facing this fate owing to human activities (IPBES, 2019; Purvis et al., 2019). In December 2022, the world’s governments adopted a new Global Biodiversity Framework, the mission of which is “To take urgent action to halt and reverse biodiversity loss to put nature on the path to recovery…” (CBD, 2022). Goal A in the framework includes commitments to halt human-driven extinctions, reduce the extinction rate and extinction risk of all species tenfold, and increase the abundance of wild species to healthy and resilient levels. Urgent conservation actions are necessary to achieve these outcomes (Leclère et al., 2020; CBD, 2022).

Crucially, we need effective conservation based on evidence (Sutherland et al., 2004), so that actions can be transparently assessed and implemented in different contexts. There is ample evidence that conservation action can work (Bolam et al., 2021; Jellesmark et al., 2022; Garnett et al., 2024; Langhammer et al., 2024) and that targeted efforts for species are needed (Bolam et al., 2023). However, this evidence is often presented in disparate sources that can be hard to access (Sutherland et al., 2019). There are also many gaps in the evidence base, including biases towards particular species (e.g. birds and mammals), geographies (e.g. North America and Europe) (Junker et al., 2020; Christie et al., 2021), and actions (e.g. protected areas) (Brooks et al., 2009). One reason for these gaps is that conservationists often lack sufficient time, capacity and/or incentives to publish outcomes of conservation actions in the scientific literature. There is a need to consolidate this information from practitioners, to understand which actions have been linked to improvements in species state, and to inform targeting of future such efforts and track progress towards halting global biodiversity loss.

The IUCN Red List of Threatened Species™ (hereafter IUCN Red List) is the most comprehensive source of information on species’ conservation (Rodrigues et al., 2006), including actions needed and underway. It incorporates input and information from a wide range of stakeholders including conservation practitioners (Mace et al., 2008). A range of information documented on the IUCN Red List is useful for assessing conservation impact, including written accounts and tabular data relating to each species’ state and the actions that are in place to promote their conservation (Salafsky et al., 2008). Although the latter can be easily analysed, some information in the narrative fields may not be reflected in the tabular fields (Challender et al., 2023; Senior et al., 2024), posing challenges for multi-species analyses, particularly those that extend beyond particular subsets of species (Smith et al., 2023; Luther et al., 2021). Correlative analyses have used these data, for example to explore the relationship between improving or worsening in species’ state or population trends and different conservation actions (Hayward, 2011; Luther et al., 2016).

Information within the Red List can also help uncover the relationships between actions implemented and species outcomes, including documenting which species have undergone genuine changes in their extinction risk, for example those resulting from increases or decreases in population size, rate of decline or distribution, rather than revisions to taxonomy or improvements in knowledge of population or range size and trends. These genuine changes underpin the Red List Index (Butchart et al., 2004; 2005; 2010), and those that have improved relate to the subset of species with the largest impacts of actions. The Red List information also allows assessments of recovery and conservation impact in the Green Status of Species fields (Akçakaya et al., 2018; IUCN, International Union for Conservation of Nature, 2021). These analyses are accessible for multiple taxa, with conservation benefit attributed to named actions. However, they rely on expert-provided information, and typically do not result from empirical tests of outcomes of actions, but instead consider relevant post-hoc assessments of counterfactual states (Hoffmann et al., 2010; Young et al., 2014; Hoffmann et al., 2015; Grace et al., 2021). However, given the species breadth and inclusion of practitioner knowledge, the different types of information in the Red List can help us gain a more comprehensive understanding of what has worked in species conservation.

We leveraged this wide range of data from the IUCN Red List to explore the following three questions: (i) which conservation actions have been implemented for different species, (ii) which species have improved in conservation state, and (iii) which actions apparently drove these improvements?

## Methods

### Data sources and handling

Information for all assessed animal species was downloaded from the IUCN Red List version 2023.1 (IUCN, 2023), including their taxonomic classification, IUCN Red List category of extinction risk, ecosystem type (terrestrial, freshwater and/or marine), threats, conservation actions in place, global population trend and generation length (in years, the average age of breeding individuals), along with their distribution (range) maps.

For comparability, only global species records were included; subspecies and subpopulations were excluded. Extinct species and taxa belonging to groups that were not comprehensively assessed (i.e. for which less than 80% of the species have Red List assessments) were excluded from analyses. We included the following groups: amphibians (7,983); birds (11,038); cartilaginous fish (1,240); cephalopods (750); dragonflies and damselflies (6,223); hagfish (80); horseshoe crabs (4); lampreys (38); lobe-finned fish (8); mammals (5,895); reptiles (10,222); freshwater fish (14,347); selected crustacea (2,886); selected gastropods (687); selected marine fish (4,988); and warm-water reef-building corals (828); for how these groups were classified see Text S1.

Generation lengths were provided as a mixture of single values or a range of values (minimum and maximum) which sometimes included a best estimate. Where available, the best estimate was used, followed by unique values. Non-numeric values were reviewed and converted to numerals accordingly (e.g. removing units, converting numbers when written as text, setting maximum and minimum values as one less or more than provided values with “<” or “>” signs respectively, etc). Any that were unclear were checked with staff in the IUCN Red List Unit. Just under half of Red List assessments for mammals were missing values, so their generation length information was supplemented from Pacifici et al. (2013). Eight species with generation lengths >150 years were checked with the IUCN Red List unit, and where incorrect, e.g. because they referred to units other than years, were corrected appropriately; only Greenland shark, *Somniosus microcephalus*, was correct.

To calculate species’ range sizes, for each species their range map with the following presence (Extant, Probably Extant or Possibly Extinct) origin (Native, Reintroduced or Assisted Colonisation) and seasonality (Resident or Breeding) codes were selected and dissolved to produce one range map per species. They were then projected into a cylindrical equal area projection before the area was calculated in m^2^. For migratory species with breeding ranges, their range size reflected their breeding (or where applicable breeding and resident) range; non-breeding ranges were excluded to avoid over-inflation of range size estimates. The area of mapped range was used rather than population size, as it was reported a greater proportion of species, with range having been shown to be correlated and so can be used as a proxy for population size (Mace et al., 2008).

Threat data were aggregated up to level 1 of the IUCN threats classification system (Salafsky et al., 2008), with the exception of Biological resource use, for which logging and wood harvesting and gathering terrestrial plants, were separated from hunting/collecting and fishing/harvesting aquatic resources, given the substantial impacts of logging and harvesting of plants on terrestrial animal species’ habitats. Future threats, or those with negligible impact, were excluded using the timing and severity coding in the IUCN threat information (threats without timing or severity coding were retained to avoid excluding important threats). To reduce the number of variables for use in modelling, threats at level 1 (except Biological resource use) were grouped into the following higher level variables: *habitat loss or degradation* (Residential and commercial development, Agriculture and aquaculture, Energy production and mining, transport and service corridors, Biological resource use: logging and wood harvesting and gathering terrestrial plants, Human intrusions and disturbance, Natural system modification and Geological Events), *hunting or fishing* (other categories of Biological resource use), *invasive or problematic species and diseases* (Invasive & problematic native species, genes and diseases), *pollution* (Pollution) and *climate change* (Climate change and severe weather).

To simplify the number of variables for analysis, where relating to the same action, the conservation actions in place were recoded to align broadly with level 1 of the IUCN Actions classification scheme. The actions in place were recoded as follows: *In protected area* (presence in at least one protected area, any threshold of protection > 0), *Area management plan* (an area-based regional management plan, typically relating to site management e.g. of a protected area), *Control of invasive or problematic species* (invasive species control/prevention), *Species management* (species action/recovery plan, species harvest management plan), *Reintroduction or translocation* (reintroduction/introduction/translocation), *Awareness and education* (recent education/awareness programmes), *Legislation or trade control* (included in international legislation, subject to international management or trade controls), *Captive breeding* or *Monitoring* (included in monitoring schemes). Note the action *Inside conservation areas* (which is intended to reflect whether sites of biodiversity importance have been identified for the species) was excluded as expert assessors’ information suggests that it has been applied inconsistently between groups, thus including uncertainty in the accuracy of this dataset.

The information on genuine changes in species Red List category was based on two sources. The first was data that underpin the Red List Index (RLI; Butchart et al., 2010; IUCN, 2023b), which includes the category change, timing, and a text narrative justifying the reason for improvement or deterioration in category (typically threats or actions). This was combined with additional genuine changes in Red List category changes identified in the period 2016 to 2022. Where a species underwent a genuine reduction in Red List category, the text explanation of the reason for the improvement was reviewed against the action codes for each species (only available for RLI data). If the specific action was not recorded (e.g. some simply listed ‘conservation action’), the species assessment and ‘actions in place’ text accounts were reviewed in the Red List to see if this contained information on clear attribution of actions to improvements in species state. Where this was absent, or an action that did not fall within the scheme was mentioned (e.g. livelihoods, economic or other incentives), they were classified as ‘other action’. When there was no clear indication that the improvement in category was driven by conservation action, or where it was attributed to natural processes (e.g. amelioration of drought, habitat succession following land abandonment, etc), this was recorded.

Information from Green Status of Species assessments (a measure of species recovery) was extracted for the 35 animal species on the IUCN Red List with published assessments (including ground-beetle, *Trechus terrabravensis*, and common birdwing, *Troides helena*, from groups that are not comprehensively assessed). These include estimates of the impact of conservation compared with a counterfactual of no action. This counterfactual is also spatially explicit, examining the impacts of conservation in distinct parts of the species range (spatial units). Within each spatial unit, the species’ current state (Absent, Present, Viable, or Functional; IUCN 2021) is estimated, as well as the counterfactual state (i.e. the expected current state without conservation action). For each species’ spatial unit, the best estimate for both the current and counterfactual state were extracted, along with documented past and current threats, and past and current conservation actions. Note this also included the action of *Livelihood, economic and other incentives* (as this is a category in the Conservation Actions Needed classification scheme but not the Actions in Place; Salafsky et al., 2008).

### Analysis of conservation actions in place

For each comprehensively assessed species group, the number of species with (i) no reported conservation action, (ii) at least one conservation action and (iii) each coded conservation action, was calculated. The list of threatened species without reported conservation actions were compared with Bolam et al. (2023)’s list of species requiring urgent management, to identify how many of these species are known to need conservation action. Chi-squared analysis and post-hoc tests were used to determine whether the proportion of species with or without any conservation action(s), and for each conservation action type in turn, varied according to IUCN Red List category and taxonomic group, and if so, how. Species assessed as Data Deficient (DD) and Extinct in the Wild (EW) were excluded because DD species lack sufficient information to assess their extinction risk, and EW species typically have captive breeding but few if any other actions in place). Species assessed as Least Concern (LC) were excluded from the chi-squared analysis across IUCN Red List categories as it is optional to report actions for LC species, so it is not possible to distinguish a lack of actions from lack of documentation.

### Models of improvements in species conservation state

To investigate which species’ characteristics are associated with improvements in species’ conservation state, three different indicators were modelled against a range of information related to species’ traits, threats and actions (summarised in Table 1). These indicators were (1) species’ global population trend (declining, stable or increasing; from species Red List assessments), (2) genuine net changes in species’ Red List category (measure of extinction risk; deteriorated, unchanged or improved), and (3) whether declines in species’ state have been prevented within specific spatial units (or not; using data from the Green Status of Species; (IUCN, International Union for Conservation of Nature, 2021).

**Table 1.** Structure of three models of improvement in conservation state against a range of variables related to species traits, threats and conservation actions. The five threats consist of: habitat loss/degradation, hunting/fishing, invasive/problematic species/diseases, pollution or climate change. The seven actions consist of: potential occurrence in protected areas, area management plans, invasive/problematic species control, species management plans, reintroduction/translocation, international legislation/trade control, education/awareness raising.

| Response variable | Model type | Fixed effects | Random effects | Taxonomic groups | Excluded cases |
| --- | --- | --- | --- | --- | --- |
| (1) Global population trend<br>*levels:*<br>*Increasing >*<br>*Stable >*<br>*Declining* | Cumulative linked mixed model | log(range) + current Red List category + system + ecosystem + 5 threat types + 7 action types | Taxonomic group | All comprehensively assessed taxa | Least Concern species, species with unknown population trends |
|  | Cumulative linked model | As above, plus log generation length for birds and mammals | N/a | Amphibians, birds, freshwater fish, mammals & reptiles separately | As above plus ecosystem excluded for except mammals |
| (2) Genuine change in Red List category<br>*levels:*<br>*Downlisted >*<br>*Stable >*<br>*Uplisted* | Cumulative linked mixed model | log(range) + initial Red List category + ecosystem + 5 threat types + 7 action types | Taxonomic group | Amphibians, birds and mammals | N/a |
|  | Cumulative linked model | As above, plus log generation length for birds and mammals | N/a | Above groups separately | Extinct in the Wild species |
| (3) Prevented declines (based on Green Status of Species)<br>*levels:* | Generalised linear mixed model (logistic) | log(generation length) + log(range) + counterfactual state + ecosystem + 5 threat types + 7 action types + livelihood/economic incentives | Species name | 35 species with published GSS assessments | N/a |
| *Prevented declines > No change in state* |  |  |  |  |  |

(1) The population trend model used data on species’ global population trend (estimated around a 5-year window of the date of assessment), excluding species with unknown population trends. This model included all species in comprehensively assessed groups, excluding Least Concern species.

(2) For the genuine change in Red List category model, the species’ Red List categories were organised by year, and species were coded as “uplisted”, “unchanged” or “downlisted” based on whether their most recent assessment category was a deterioration, the same, or an improvement compared to their initial category. The model was restricted to amphibians, birds and mammals, as these are the only comprehensively reassessed groups with sufficient numbers of species that improved or deteriorated in Red List category and that have been assessed more than once.

(3) For the model of prevented declines in species state within specific spatial units, the changes in species state between the current and the counterfactual state (without conservation action) were documented for each spatial unit. If there was an improvement in state, this was coded as a ‘prevented decline’, and otherwise coded as ‘no impact’. No spatial unit had a worse current state compared to the counterfactual state. Species’ spatial units that were classified as “functional” in both current and counterfactual states in the analysis were excluded (as they could not improve). This model only includes the 35 animal species with published Green Status of Species assessments.

The three indicators of improvements in species’ state were modelled against a range of covariates. Species’ range size (log m^2^) was included as this has been shown to be a key indicator of extinction risk (Purvis et al., 2000; Mace et al., 2008). Species’ ecosystem type (terrestrial, marine or both terrestrial and marine) was included to account for differences in threats and actions between the different realms; with freshwater included in terrestrial to focusing on the distinction between land and sea where species traits, threats and actions are likely to differ more substantially. The other factors included in the model were the presence/absence of the 5 threats and 7 actions listed above. The population trend model was also run with only species traits, so Least Concern species could also be included. Log of generation length (another important indicator of extinction risk; Mace et al., 2008) was explored but was not available for many species, so it was included within models of: species characteristics only, birds and mammals, and the prevented declines in species state within spatial units.

Species’ current IUCN Red List category (for the population trend model) or initial Red List category (for the genuine Red List category changes model) was also included as an ordered factor, where Least Concern < Near Threatened < Vulnerable < Endangered < Critically Endangered < Extinct in the Wild. Least Concern species were included in the genuine Red List category change and prevented decline in state models, as threat and action information for these subsets of species were more likely to be complete to evidence genuine changes and given species would have moved into or out of threatened or Near Threatened categories, and to support Green Status of Species assessments respectively. The prevented declines in state model included the counterfactual state within each spatial unit, and an additional action of *Livelihood, economic & other incentive*s included in the prevented declines model. Note the inclusion of the species in one or more monitoring schemes was excluded from the models (not in other parts of analysis) as these relate to research and monitoring rather than conservation interventions per se. Ex situ conservation (captive breeding) was excluded from the models as it does not have an impact on species’ extinction risk until individuals are introduced into the wild (which is documented separately as ‘species reintroduction’ or ‘benign introduction’). Taxonomic group was included as a random factor in the models for population trend and genuine Red List category changes. Due to the limited number of species in the Green Status of Species model, species name was included as a random factor rather than taxonomic group. Including species name as a random factor made sense within the Green Status of Species model, because the assessment divides the species into multiple spatial units, and so there were often multiple observations for a species in the model.

Cumulative linked mixed models were used to model the population trend and genuine change in extinction risk, and generalised linear mixed models (with a logistic function) were used to model prevented declines in species’ state in spatial units. Backward selection was used to find the best model, dropping the variable with the highest p-value > 0.05 in turn until all remaining variables were significant (Bruce et al., 2020). Models were run across all species within the respective dataset, and species group models were run separately for both population and Red List category change models, where species had sufficient sample sizes to converge (see Table 1). The prevented declines model did not have sufficient numbers of species in any one group to evaluate independently. Model estimates were then visualised in a heatmap to enable visual comparison of the relationships of different variables with the different indicators of improvement in state within and between the various models.

To explore how many species have undergone net movement between each Red List category, the numbers of species moving from and to each category was also visualised in a contingency table. To visualise the distribution of species that underwent genuine changes in Red List category, their ranges were loaded into QGIS Desktop version 3.22.12 (QGIS, 2024). Ranges of species that underwent genuine net improvement or deterioration in Red List category were then in turn spatially joined to a fishnet grid (at 1 decimal degree resolution), to count the number of species per grid cell that improved or worsened in category.

### Actions driving genuine changes in Red List category

For species that underwent genuine changes in Red List category, we calculated the number of species with (i) no action in place, (ii) at least one action in place, (iii) each of the possible actions in place (excluding the ‘other conservation actions’), (iv) each action in place that was judged by assessors as driving the genuine change in Red List category. This was done for birds and mammals independently, as these were the only groups with data on genuine changes in Red List category, and with sufficient numbers of species having a documented reduction in extinction risk with actions judged by assessors as driving genuine improvement. Chi-squared and post-hoc tests were used to evaluate whether there were differences in the types of actions that led to improvements relative to those in place. This was analysed for both birds and mammals combined, and independently.

### Software

All analysis was undertaken in RStudio (Posit team, 2024) using R version 4.3.1 (R Core team, 2023). This included use of the following packages for data manipulation: tidyverse (Wickham et al., 2019); statistical analysis: chisq.posthoc.test (Ebbert, 2019), lme4 (Bates et al., 2015), ordinal (Christensen, 2023), ggcorrplot (Kassambara & Patil, 2023); plotting: ggplot2 (Wickham, 2016), pheatmap (Kolde, 2019); and spatial analysis: sf (Pebesma, 2018; Pebesma & Bivand, 2023).

## Results

### Numbers of conservation actions in place for species

Over half of species (34,847 out of 67,217; 51.8%) in comprehensively assessed animal groups had actions in place documented, rising to 58.7% (7,380 of 12,574) for threatened species. Across all assessed species, just under half of species had at least one reported conservation action (42,980 of 89,076 assessed species, 48.3%), with a slightly higher proportion for threatened animal species (9,068 of 17,416, 52.1%). Of the 5,194 threatened species (1,324 of which are Critically Endangered) that do not have any conservation actions reported; 41.0% are freshwater fish (2,132), 13.7% are amphibians (710) and 12.6% are reptiles (656), with 29.3% (1,523), 21.1% (1,095) and 18.4% (954) of these species without actions reported found only in the Neotropical, Indomalayan or Afrotropical realms respectively. We found at least 778 of these species required urgent management action (based on Bolam et al., 2023’s data).

Among comprehensively assessed taxa, warm-water reef-building corals and terrestrial tetrapods were more likely to have at least one conservation action reported compared with other invertebrates or aquatic species (Figure 1A; Table S1; X^2^ = 15364, df = 12, p < 0.001). Near Threatened or Extinct in the Wild species were more likely to have conservation actions reported compared with threatened species (VU, EN, CR) or Data Deficient species (Figure 1B), with Critically Endangered species least likely to have actions and Near Threatened species most likely (Table S1; X^2^ = 179.1, df = 3, p < 0.001). However, this pattern varied across taxonomic groups, with more highly threatened amphibians, reptiles, dragonflies and damselflies, and freshwater fish less likely to have actions in place than less threatened species, but birds, mammals, crustacea, marine and cartilaginous fish showing the opposite pattern (Figure S1).

**Figure 1.**
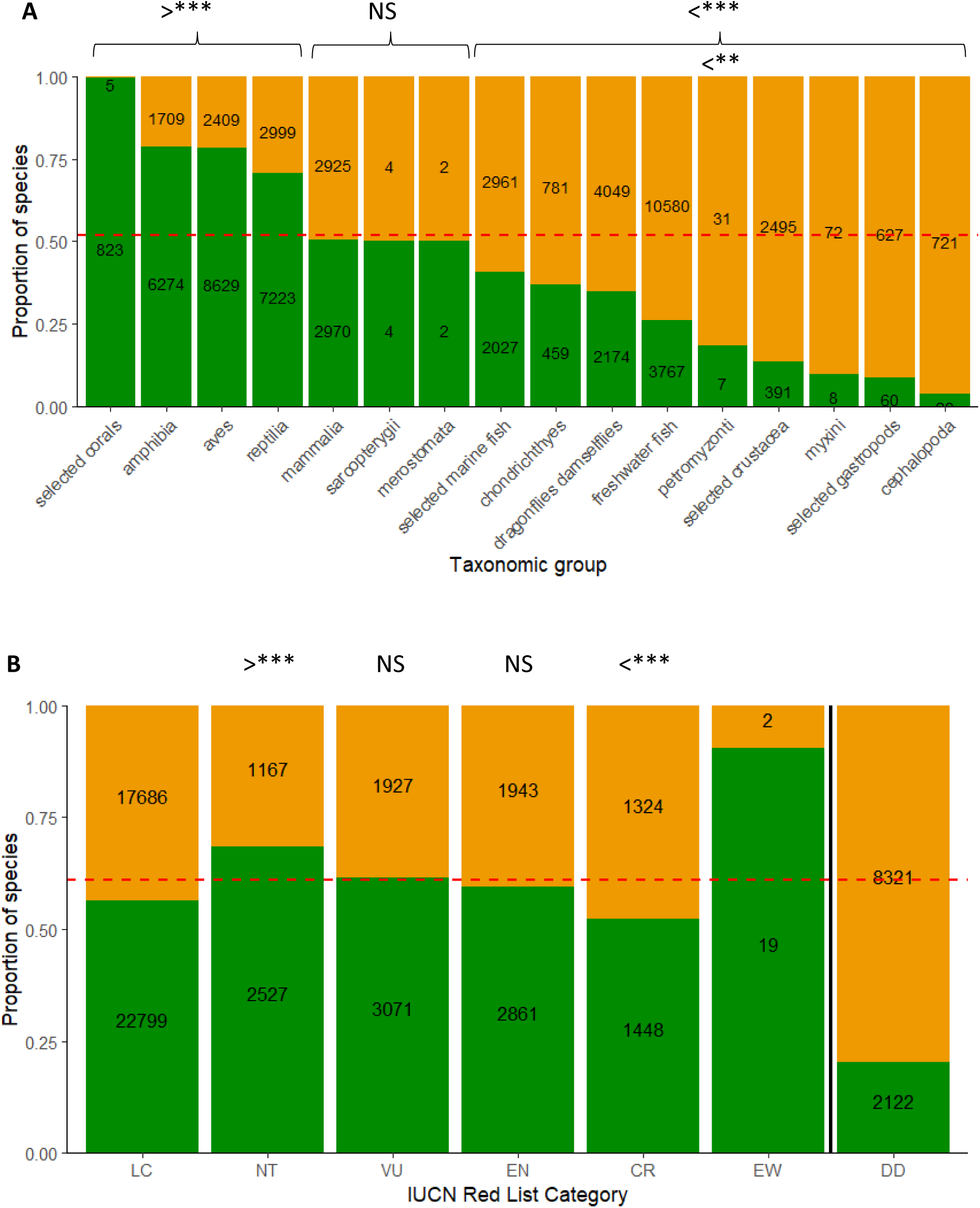
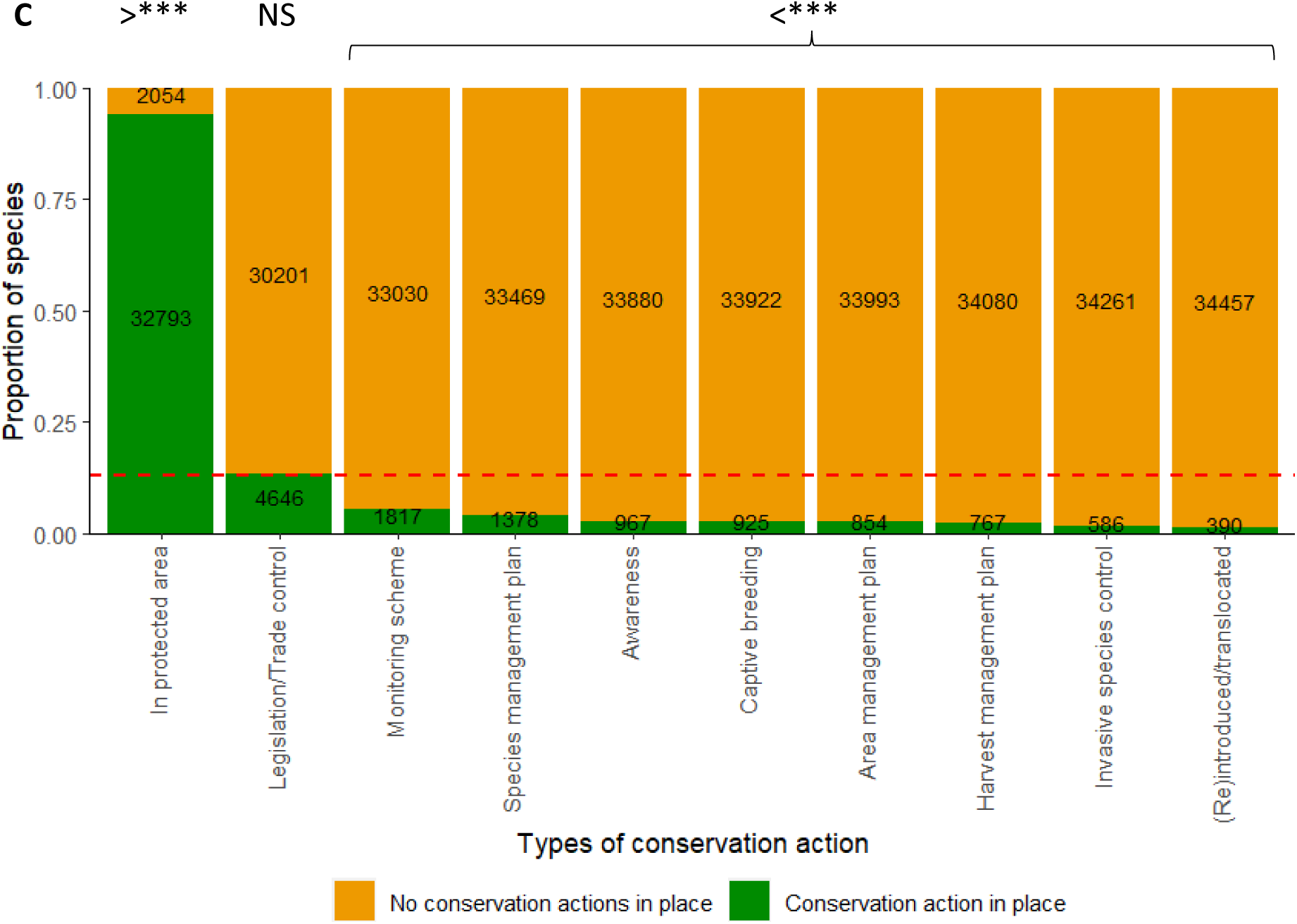
The proportion of species (in comprehensively assessed taxonomic groups) with at least one conservation action documented as being in place, for (A) different taxonomic groups, (B) different Red list categories of extinction risk, and (C) for different conservation actions. The dashed orange line indicates the mean proportion across all species (in B; LC, EW and DD were not evaluated). The > and < signs indicating significantly more or less likely to be in place compared to other actions (evaluated using a chi-squared test in Table S1), with statistical significance level indicated as follows: p < 0.01 **, p < 0.001 *** and p > 0.05 NS.

### Types of conservation actions in place for species

For species with conservation actions reported, the most common action reported across comprehensively assessed groups was potential occurrence in at least one protected area (32,793 of 34,847, 82.7%; Figures 1C and S2A; Table S2; X^2^ = 229,674, df = 9, p < 0.001). Along with protected areas, birds and mammals were more likely to have legislation or trade control and cartilaginous fish and crustaceans were more likely to have harvest management in place compared to other action types respectively. Tetrapods, particularly birds and mammals, report a wider variety of actions, compared to aquatic and invertebrate groups that mostly only report potential occurrence in protected areas (Figure S2A).

Species with conservation actions reported and at a greater threat of extinction were more likely to have a range of different types of species-targeted conservation actions in place (compared to broader actions like protected areas), with species at lower risk of extinction being more likely to be reported to potentially occur within protected areas than species at higher risk (Table S2; Figure S2B; X^2^ = 513.9, df = 27, p < 0.001). Near threatened species were more likely to have occurrence in protected areas, and less likely to have captive breeding, species management plans or awareness or education actions in place. Vulnerable species were also less likely to have awareness or education actions in place. Critically Endangered species were more likely to have captive breeding, species management plans, monitoring schemes and invasive species control in place.

### Species with improvements in conservation state

Although few species in comprehensively assessed groups have increasing global population trends (969 of 67,217 species, 1.4%), the majority of these have at least one conservation action in place (759 of 969 species or 78.3%). Since 1980, 289 species are reported to have undergone genuine improvements in their Red List category through conservation or other reasons (i.e. to have qualified for downlisting), consisting of 122 amphibians (since 1988), 111 birds (since 1988), 33 mammals (since 1996), 2 warm-water reef-forming corals (since 1996), and 6 reptiles, 6 marine fish, 4 freshwater fish, 3 cartilaginous fish and 2 snails (all since 2016). Of the 289 species that underwent genuine reductions in extinction risk, only 13 species had no reported actions in place (six amphibians, four birds, two reptiles and one mammal), of which 11 had conservation actions indicated in the narrative text, leaving only two species with no apparent actions in place, *Craugastor taurus* (amphibian) and *Notomys cervinus* (mammal). Considering only amphibians, birds and mammals, far more species underwent net deterioration (1,220) in Red List category than net improvement (202), with most species moving one category level within the past 44 years (Table 2). Although 25 species moved from Least Concern to Critically Endangered, no species moved from Critically Endangered to Least Concern.

**Table 2.** The total number of species which started and ended in each respective combination of IUCN Red List categories between 1980 and 2024. Net genuine improvements in category are written in green, deteriorations in orange, and unchanged in black.

|  |  | Previous Red List Category |  |  |  |  |  |
| --- | --- | --- | --- | --- | --- | --- | --- |
|  |  | LC | NT | VU | EN | CR | EW |
| Current Red List Category | LC | 15711 | 25 | 8 | 1 | 0 | 0 |
|  | NT | 336 | 1416 | 46 | 30 | 10 | 0 |
|  | VU | 144 | 149 | 1738 | 30 | 15 | 0 |
|  | EN | 45 | 50 | 146 | 1913 | 34 | 1 |
|  | CR | 25 | 21 | 92 | 207 | 925 | 2 |
|  | EW | 0 | 0 | 0 | 0 | 5 | 3 |

Amphibians, birds and mammals that underwent genuine deterioration in Red List category were found across the tropics, southern Europe, parts of central Asia and south-eastern Australia, with highest concentrations in the tropical Andes, Peninsula Malaysia, Sumatra and Borneo (Figure 2A). Species that underwent genuine improvements in Red List category were less concentrated, with the highest numbers of species on islands, such as New Zealand, Mauritius, the Seychelles, Chatham Island, Guadeloupe and the northern tip of Borneo, as well as in parts of the eastern United States of America, Costa Rica, eastern Australia, the southern tip of India and isolated patches in Mexico, Ecuador and Brazil’s Atlantic Forest (Figure 2B). Note coastline numbers are slightly inflated due to overlaps with several seabird and whale species’ ranges.

**Figure 2.**
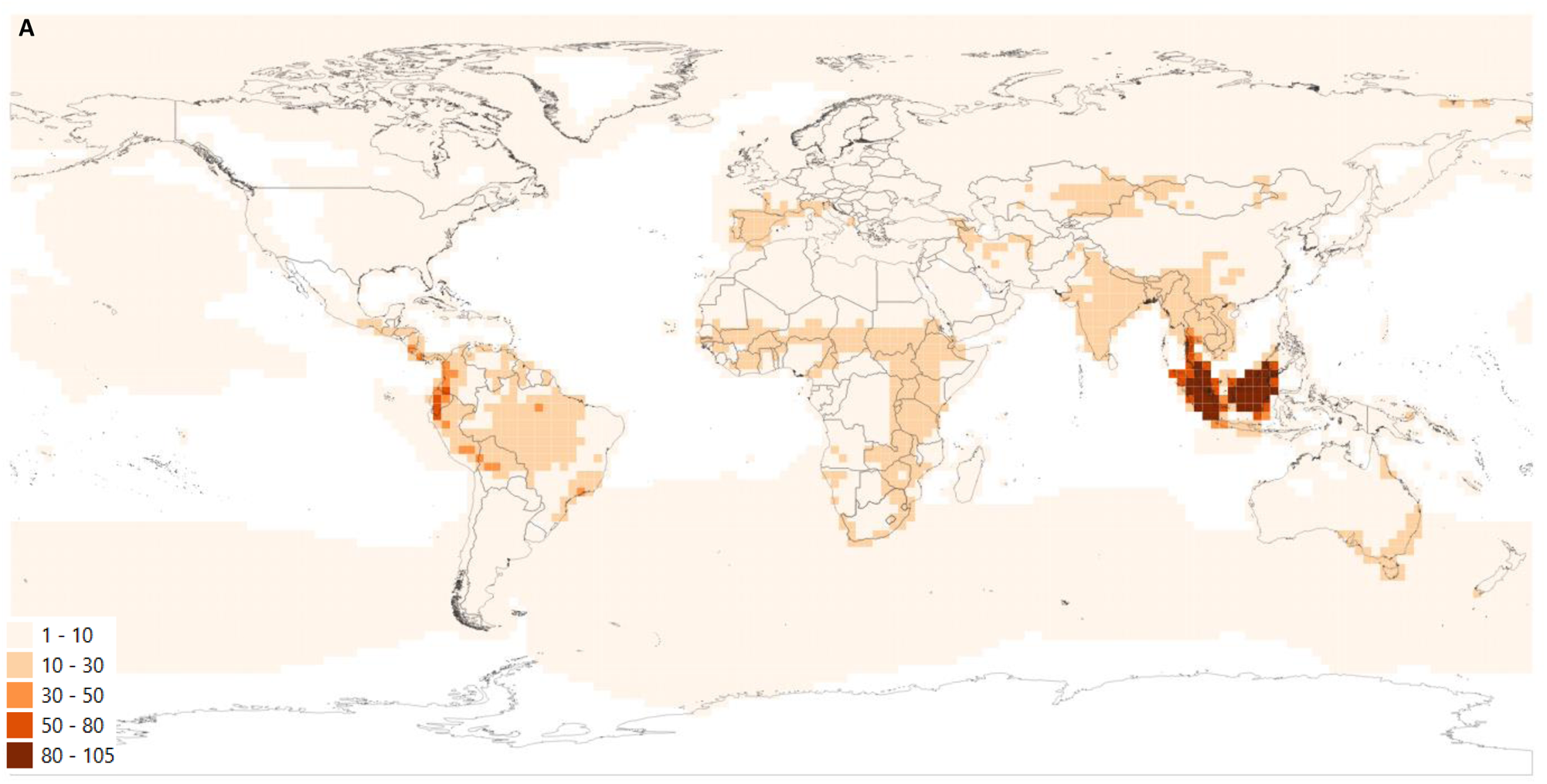
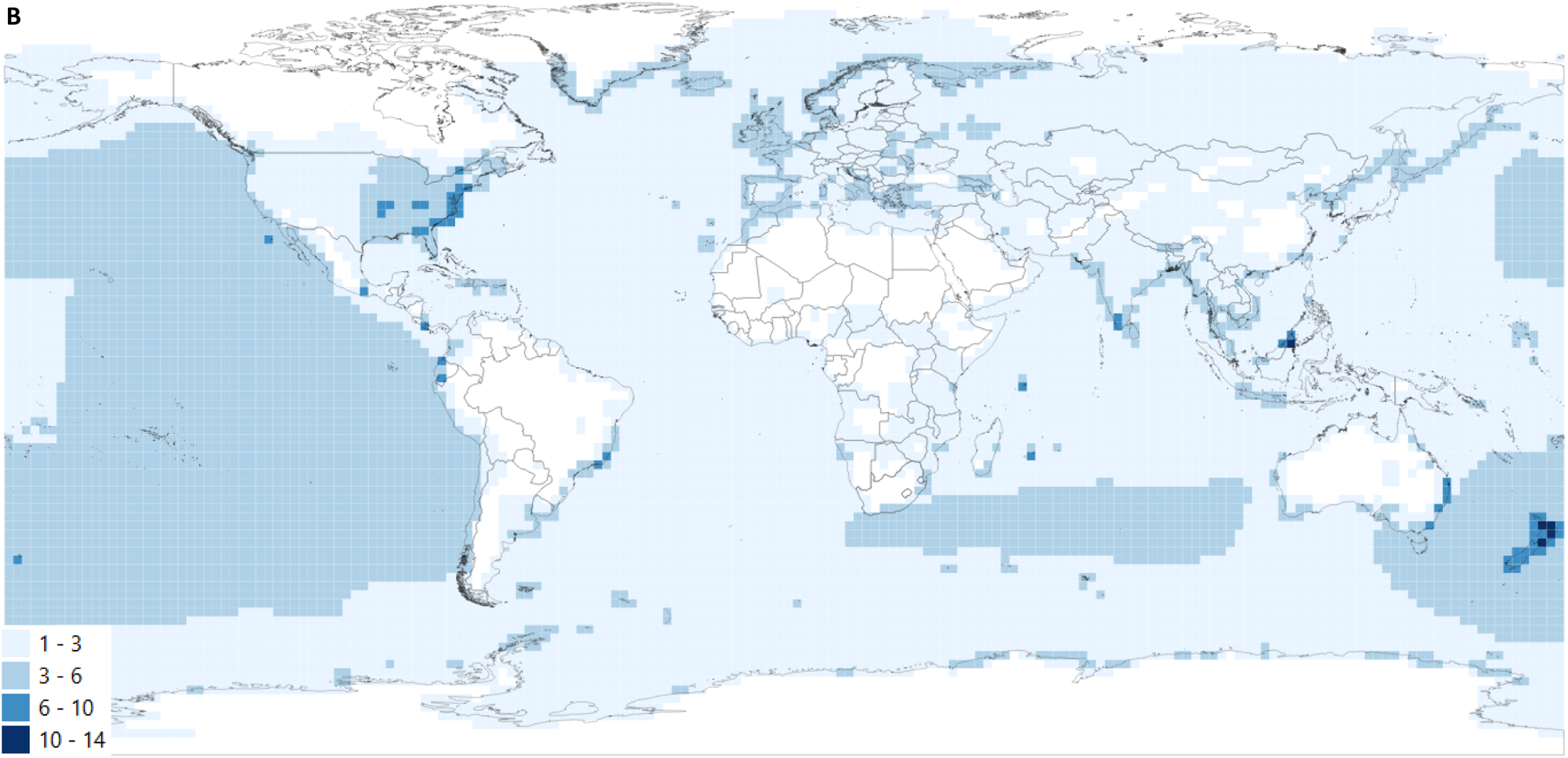
Species richness map of species that have undergone genuine deteriorations (orange; A) or genuine improvements (blue; B) in Red List category (measure of extinction risk). Lighter colours indicate fewer species and darker colours indicator high numbers of species. White indicates areas where no species underwent a change in Red List category.

### Species traits

Species with smaller global ranges and those at lower risk of extinction were more likely to have increasing global populations (Figure 3; Table S3). By contrast, species with larger global ranges and previously at higher risk of extinction were more likely to have improved in Red List category (extinction risk). For species with Green Status of Species assessments, species that would be absent without conservation are most likely to have had prevented declines in their state within spatial units, compared to those that would be in more favourable states. Species present in both terrestrial and marine ecosystems were more likely to have increasing population trends (such as species of seals and seabirds), and those restricted to marine ecosystems were more likely to have undergone genuine improvements in Red List category, compared with entirely terrestrial species (Figure 3; Table S3). Species with shorter generation times were more likely to have increasing global population trends and have improved in Red List category, except for birds and mammals, for which species with longer generation times are more likely to have increasing global populations (Figure 3; Table S3).

**Figure 3:**
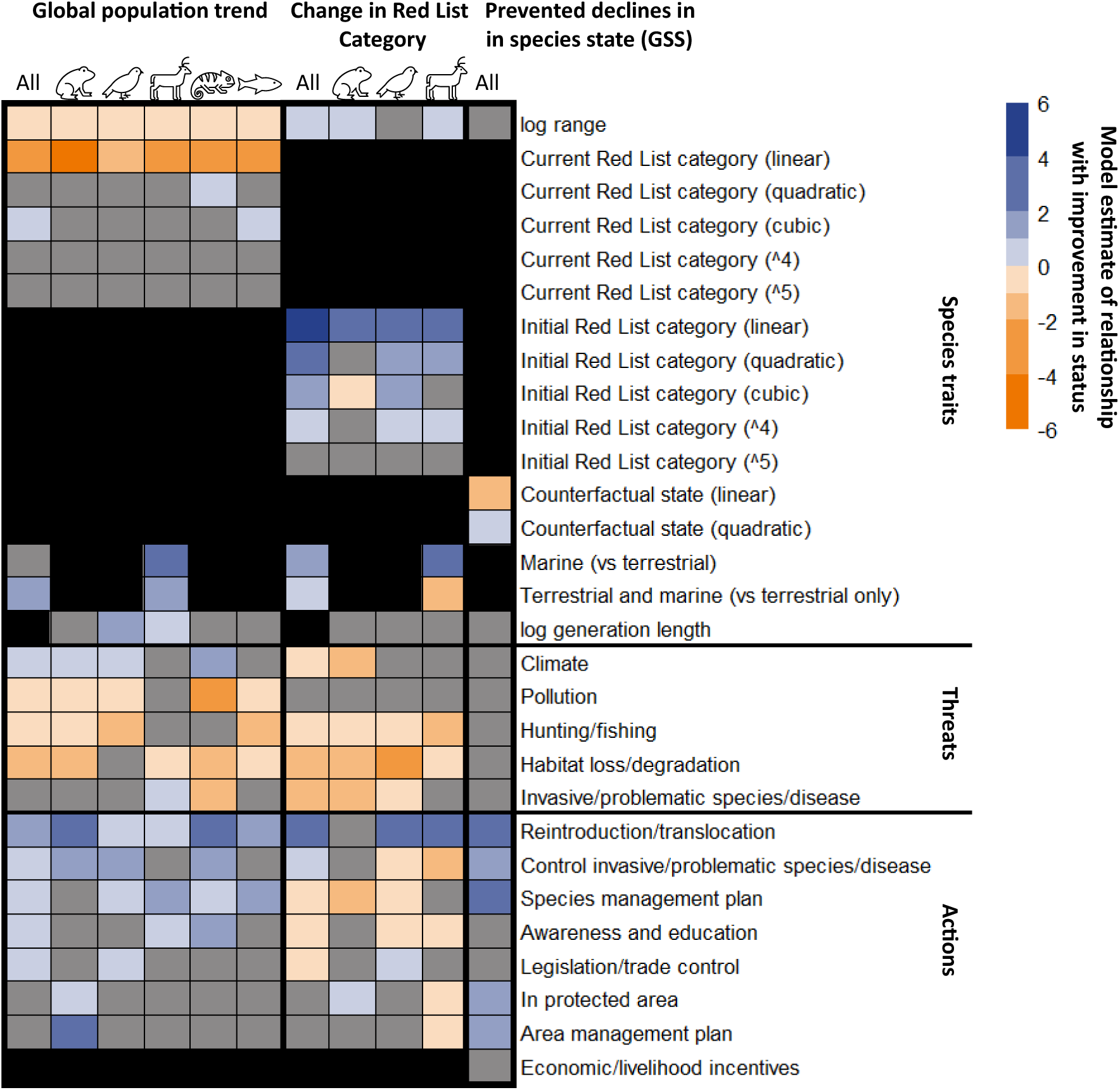
Estimated relationship between various species traits, threats and actions in place with three indicators of improvement in species state; species’ global population trends, genuine changes in Red List category (both derived from cumulative-linked mixed models) and prevented declines in species’ state in spatial units (from the Green Status of Species; derived from a Generalised Linear Model). Modelled outputs are shown across all species and for different groups of species in turn (amphibians, birds, mammals, reptiles and freshwater fish). Black indicates variables that were not included in the initial model, and grey indicates variables that were dropped through backwards selection. Darker colours indicate largest variable estimates, with blue indicating larger positive associations and orange larger negative associations with improvements in species state. See Table S3 for full model outputs.

### Threats

Species threatened by habitat loss or degradation and hunting or fishing were more likely to be undergoing population declines or worsening in Red List category. Species threatened by pollution, problematic species, disease or climate change, were more likely to have declining populations or to have deteriorated in Red List category respectively. By contrast, species threatened by climate change and mammalian species threatened by problematic or invasive species or diseases were more likely to have increasing populations.

### Actions

Species that were reintroduced or translocated, or have invasive or problematic species or disease control in place were more likely to have improved in conservation state across all three indicators (Figure 3; Table S3). Species with species management plans were more likely to have increasing global populations and have had prevented declines in state within spatial units (Green Status of Species), whilst species with education or awareness raising actions or international legislation or trade control, and area management plans and potential occurrence in protected areas, were more likely to have increasing populations or have had prevented declines in state within spatial units respectively.

For amphibians, species potentially occurring in protected areas were more likely to have increasing population trends and have undergone improvements in Red List category, and those with area management plans were also more likely to have increasing populations. Birds with international trade or legislation in place were more likely to undergo improvements in Red List category (Figure 3; Table S3). By contrast, overall, species with international legislation or trade control, awareness or education actions, or species management plans in place were more likely to have deteriorated in Red List category. Birds with control of invasive or problematic species or disease in place and mammals with control of invasive or problematic species or disease in place, potentially occurring in protected areas and area management plans in place were more likely to have deteriorated in Red List category.

### Drivers of genuine improvements in Red List category

Improvement in Red List category was attributed to conservation action in 91 species (71% of 128 species of mammal and birds), comprising of 75 bird and 16 mammal species; of which 63.7% (58) of species had two or more conservation actions attributed and 36.3% (33) had one action attributed. Only seven bird species, such as Woodford’s rail, *Hypotaenidia woodfordi*, and Mewing kingfisher, *Todiramphus ruficollaris*, had only non-conservation reasons for improvement, relating to natural recovery following severe weather events, land abandonment or cessation of other activities. For the remaining 24 bird and 6 mammal species, the reason for improvement was not clearly attributed.

Across birds and mammals, there was a significant difference between actions in place across groups compared with those that assessors determined to have driven the reduction in extinction risk (X^2^ = 71.739, df = 9, p < 0.001). Reintroductions or translocations were significantly more likely to have led to reductions in extinction risk compared with other actions, and monitoring activities were less likely to have led to reductions in extinction risk (Figure 3; Table S3).

For birds that improved in Red List category, potential occurrence in protected areas (86) was the most reported action, followed by species having a management plan (69). However, area-based regional management plans and reintroduction/translocation (both 30) and invasive or problematic species or disease control (27) were the most commonly attributed actions, with legislation and trade control (6) and monitoring schemes (3) less likely to be attributed (X^2^ = 75.418, df = 9, p < 0.001; Figure 4A; Table S4).

**Figure 4.**
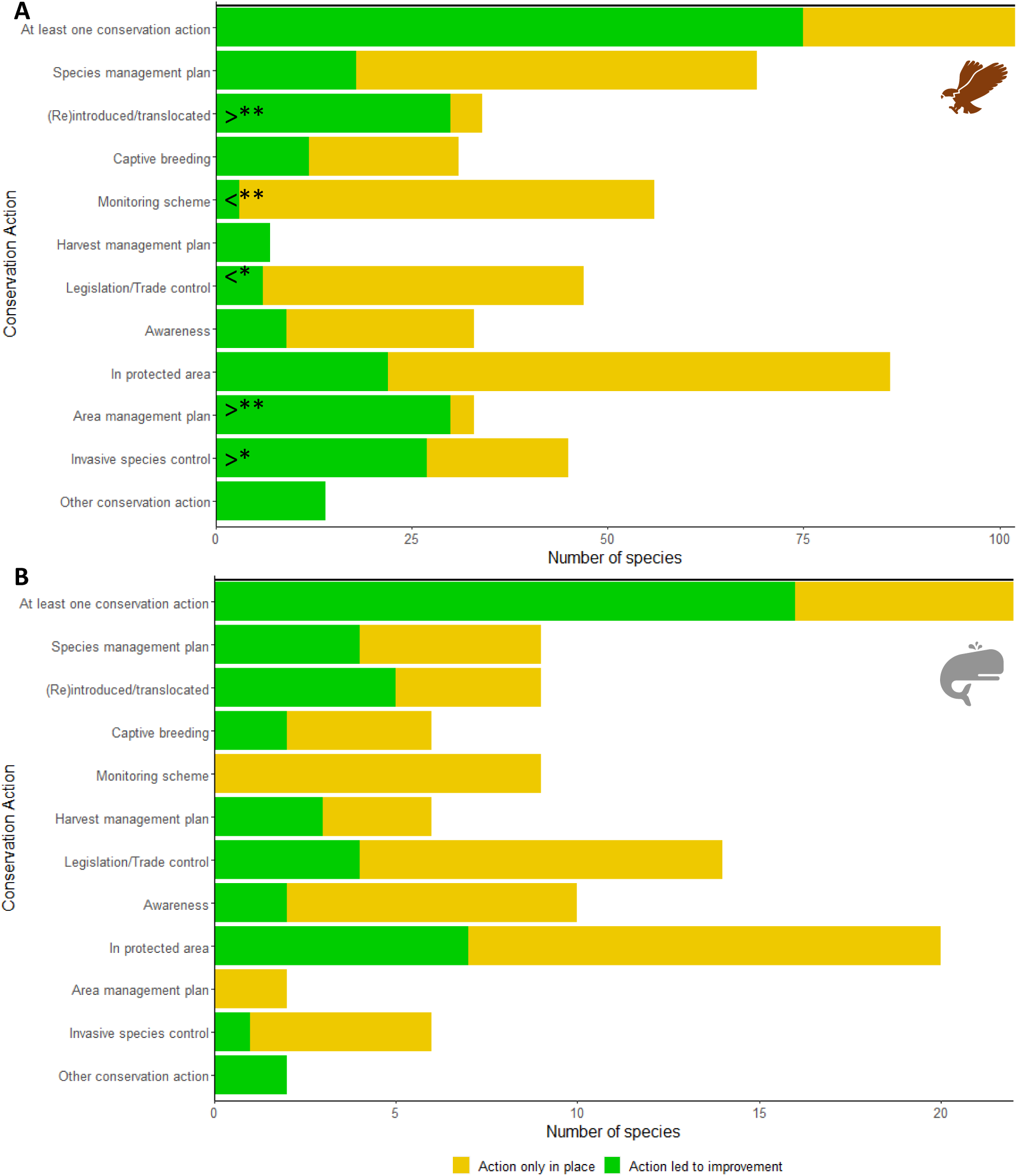
Actions in place for (A) birds and (B) mammals that have shown a genuine improvement in Red List category (and that have at least one conservation action in place), and whether any of those actions were deemed by experts to have contributed towards the genuine improvement. Asterisks indicate where actions were responsible for improvements in Red List category more than expected (determined by chi-squared analysis; Table S4).

For mammals that improved in Red List category, the most common actions were potential occurrence in protected areas (20) and legislation/trade control (14). Potential occurrence in protected areas (7) and reintroduction or translocation (5) were the most frequently attributed actions, and reintroductions or translocations, harvest management and species management most proportionately attributed (relative to how often they are in place), although a significant difference in attribution across the different action types could not be detected (X^2^ = 9.509, df = 7, p = 0.218; Figure 4B).

## Discussion

### Which conservation actions have been implemented for different species?

Just over half of all species in comprehensively assessed animal groups, and almost 60% of threatened species, have documented conservation actions in place, a slight increase of estimates from Senior et al. (2024), likely reflecting updates to species assessments and inclusion of additional animal groups. Terrestrial tetrapods, warm-water reef-building corals and species at low risk of extinction were more likely to have actions reported compared to fish, aquatic invertebrates and taxa at high risk of extinction. The suggested greater focus on terrestrial tetrapods is hardly surprising and supports findings by many studies including those evaluated by Di Marco et al. (2017). Corals are an exception, as they form the basis of an ecosystem and so may be more likely to be within protected areas.

Protected areas were the most frequently reported action across species. Protected areas also likely explain why Near Threatened species were more likely to report actions than more at risk species, as Near Threatened species typically have larger ranges and are therefore more likely to overlap with at least one protected area, and these species were much less likely to have other action types reported. Although Critically Endangered species were less likely to report actions, those with actions were more likely to have different species-targeted actions in place (such as species management and captive breeding) compared to more general actions such as protected areas, possibly reflecting greater targeting and or documenting of conservation efforts to species most urgently requiring them (Luther et al., 2016; Senior et al., 2024). Comprehensively assessed groups of crustacea and cartilaginous fish were more likely than other species groups to have harvest management actions in place, presumably because they are particularly targeted by fishing and/or threatened by bycatch.

Given that reporting actions in place is only recommended in the Red List, the estimated number of species with actions likely represents an underestimate, particularly for Least Concern species where it is optional. It is likely that some of the species have information in the Conservation Action narrative that is not reflected in the tabular data (Challender et al., 2023; Senior et al., 2024). Reported actions across taxa may also differ between Red List authorities and assessors; all bird assessments are undertaken by BirdLife International (as the designated Red List Authority for birds), and they have been assessed eight times since 1988, so the documentation is likely to be more consistent and comprehensive. By contrast, mammals have only been comprehensively assessed twice since 1996 by over 35 different groups IUCN Species Specialist Groups, with more than 100 of these groups existing across all assessed taxa (SSGs), so greater variation in documentation and approach might be expected. Also some actions, such as protected areas and international legislation (UNEP-WCMC & IUCN 2024; (*CITES Trade Database*, 2024), are likely disproportionately represented due to the availability of global databases, enabling fairly rapid assessment across a large number of species.

However, the number of species with actions underway arguably exaggerates the degree to which species have been specifically targeted for action, particularly for protected areas (Senior et al., 2024). While some protected areas were designated and/or are managed to conserve particular species, the many are not and instead are aiming to protect an important area and its broad species assemblage (Dudley, 2008; Geldmann et al., 2013). Furthermore, given the large coverage of the world by protected areas (14.6% of land and 2.8% of marine areas), it is likely many species’ ranges overlap protected areas only marginally (S. H. M. Butchart et al., 2015), with recent studies showing around 91% of threatened species have insufficient representation of their habitats within protected areas (Senior et al., 2024). Those protected areas that do overlap ranges may also be located in less accessible areas, under lower human-pressure and so not safeguard species from threats (Joppa & Pfaff, 2009; Rodrigues & Cazalis, 2020), or may lack effective management (Wauchope et al., 2022).

Despite gaps in action reporting, we should not ignore the 5,194 threatened species (41.3% of the 12,574 threatened species) with no reported actions in place. At least 778 were also identified by Bolam et al. (2023) as needing urgent management actions, including the Critically Endangered Estuarine Pipefish, *Syngnathus watermeyeri,* and the Endangered Mount Namuli Pygmy Chameleon, *Rhampholeon tilburyi*. The 4,416 other threatened species are priorities to have their actions assessed, and if there are genuinely no actions in place such as the Critically Endangered Yellowmargin Gudgeon, *Allomogurnda flavimarginata*, they should be urgent priorities for conservation attention and action. We highlight the importance of documentation of actions underway for all species when undertaking Red List assessments, alongside the extent and impact of actions, to better identify conservation priorities.

### How are species changing in Red List category?

Since 1980, six times more species have deteriorated rather than improved in Red List category. Threats have driven species from the lowest to highest level of extinction risk, but few actions have reversed this trend. Large numbers of species that deteriorated in Red List category were found across the land surface, with very high concentrations in biodiversity hotspots, mostly driven by habitat loss or degradation or hunting, such as the Binturong, *Arctictis binturong,* in Borneo. By contrast, species that improved in Red List category were less concentrated, with largest concentrations typically found on islands or in localised habitat patches; examples include the Pink Pigeon, *Nesoenas mayeri*, in Mauritius. This suggests broad-scale threats such as habitat loss and degradation are driving more widespread declines of species (Hayward, 2011), with limited success of spatially localised actions to recover species to resilient levels.

### Which species have improved in conservation state?

#### Species traits

Species with smaller ranges, typically found in isolated habitat patches or on islands, were more likely to have improved in conservation state, perhaps as conservation challenges are more circumscribed. Although Red List category appeared to indicate the opposite, this likely reflects modelling of current rather than past range size, which may have increased following improvement in category (compared to reductions in range size for those that deteriorated), and range size information being partly captured by the Red List category. Species that were entirely marine or occur in both terrestrial and marine ecosystems were more likely to have improved in state than entirely terrestrial species, perhaps due to less concentrated area-competition with people compared to on land. Species previously at higher risk of extinction were more likely to have undergone a genuine reduction in extinction risk, which may relate to both greater targeting of conservation efforts towards species most at risk (D. A. Luther et al., 2016) and the greater potential to increase population numbers above thresholds needed for downlisting at lower levels of population size (where thresholds are lower). This means it is easier for a species to pass from Critically Endangered to Endangered than from Near Threatened to Least Concern.

Species with shorter generation times were more likely to improve in state, possibly because a faster rate of reproduction can enable more rapid recovery of their populations. Bird and mammal species showed the reverse relationship, potentially because such species are typically larger and therefore tend to receive more conservation attention and resources (Berti et al., 2020; Malhi et al., 2022). Alternatively, many of these species may already have largely depleted populations and so are increasing from extremely low levels (consistent with Red List category result), such as the European bison, *Bison bonasus,* which is currently undergoing a spectacular recovery from having been driven to extinction in the wild by hunting and habitat loss, yet still only occupies a small fraction of its former range (Plumb et al., 2020).

#### Threats

Unsurprisingly, species impacted by any of the five major threats to biodiversity (habitat loss and degradation, hunting or fishing, invasive or problematic species or diseases, pollution and climate change) were more likely to have declined in population trend or deteriorated in Red List category, with habitat loss and degradation, and hunting and fishing, indicating deteriorations in both indicators. Where species with a reported threats showed improving state, this likely reflects issues related to the reported threat data. This could relate to the inclusion of threats in the models where the impact or extent of the threat on some species was reported as unknown, may reflect a limited number of species involved, or the actions that were in place may have been able to mitigate or offset the impacts of the threat.

### Which actions were associated with or attributed to improvements in species’ state

Almost all species with increasing global population trends or that experienced a genuine reduction in their extinction risk had at least one conservation action in place, with the majority of bird and mammal species that improved in state due to conservation action. By contrast, Luedke et al. (2023) found roughly half of amphibian species improved due to conservation action and the other half from a natural capacity to recover from chytridiomycosis. In our analyses, species with any of the broad types of conservation action in place were more likely to have improved in state in at least one of the models, with each of these actions attributed by experts to the improvement of at least one species. This reinforces similar findings that most conservation action works (Hoffmann et al., 2010; Langhammer et al., 2024).

Reintroductions or translocations, and the control of invasive or problematic species or diseases were consistently associated with positive indicators of species conservation state across species, and significantly attributed to improvements of bird species. Reintroductions or translocations, when successful, will significantly increase the wild population or distribution of a species and therefore reduce extinction risk (*IUCN Red List Categories and Criteria, Version 3.1, Second Edition*, 2012; Mace et al., 2008). For example, the intensive captive breeding and reintroduction lead to the recovery of the Mauritius Kestrel, *Falco punctatus,* with numbers in the wild increasing from 4 wild birds to >250 mature individuals and subsequent downlisting from Critically Endangered to Vulnerable (BirdLife International, 2023; though has since been uplisted to Endangered due to subsequent declines). However, reintroductions can also fail; this can be due to the way they are implemented or failure to remove the original driver of extinction (e.g. hunting and extirpation of Arabian oryx, *Oryx leucoryx*, following reintroduction in Oman; Spalton et al., 1999). Reintroductions in particular, but also many other actions, are often only successful alongside other actions; our findings supported this with almost two thirds of species which improved in Red List category having two or more actions attributed. An example includes the conservation of the Western Quoll, *Dasyurus geoffroii,* which is thought to have improved in Red List category as a result of translocation, invasive predator (feral cats) control, and public awareness raising (Woinarski & Burbidge, 2019).

Similarly, the positive impact of invasive control is consistent with the many reported successful control programmes, particularly on islands (Jones et al., 2016), such as successful eradication of invasive rats from Campbell Island for the recovery of Campbell Teal, *Anas nesiotis* (BirdLife International, 2020). However, these also fail, such as the mouse eradication on Gough Island (*The Gough Island Restoration Programme*, 2022) or rat eradication on Henderson Island (Amos et al., 2016), or may not be carried out sufficiently across species’ ranges, perhaps explaining the association with worsening in state of some bird and mammal species.

Similar to Luedtke et al. (2023), we found amphibians appeared to benefit from area related actions, particularly area management plans and potential occurrence in protected areas, perhaps enabled by their typically smaller ranges compared to other terrestrial vertebrates which makes it easier to target such interventions by including a larger percentage of their range in the protected area. Experts also attributed area management to the improvement of many bird species, reinforcing past findings (D. A. Luther et al., 2016), such as the Dark-tailed Laurel-pigeon, *Columba bollii*, which recovered from Near Threatened to Least Concern due to increased protection and restoration of native laurel forests in the Canary Islands (BirdLife International, 2016). Management of important sites for species of conservation concern (e.g. Important Bird and Biodiversity Areas and other Key Biodiversity Areas), whether through formal protected areas, other effective area-based conservation measures, or other means, can generate substantial benefits for species for which site-scale conservation is appropriate (S. H. M. Butchart et al., 2012; Pavón-Jordán et al., 2020).

### Actions with mixed associations with species state

Change in Red List category was the only indicator with negative associations with actions, both across and for particular taxa. Where actions were either not significantly associated or negatively associated with one of the indicators, this may be because either actions were not carried out at sufficient scale to impact the species’ global Red List category or trends, other threats may be acting which the action did not address, a lag in species responses, declines slowed but not yet halted, reversed or sufficient to change Red List category, or the species has not been reassessed. Evaluations of conservation success can thus represent important opportunities to consider such aspects and the need to implement other or additional actions.

Model disagreement may also arise from the variation in species and species groups included, the indicator resolution, and the temporal and spatial scales of the data. Both the population trend and Red List category models analyse changes to species’ global state, but with the population model considering a wider range of species and species groups but with a broader indicator of improvement, where the Red List category allow a more nuanced measure of state change. By contrast, the prevented decline data only considers 35 species with a coarse expert derived counterfactual category of improved or not, but examines spatial units within species’ ranges, covering time between present and as far back as 1500. This data from the Green Status of Species can therefore identify changes to species’ states for different populations of a species and changes within Red List category levels, including slowed decline rather than just improvements in state, alongside impacts of conservation action further in the past.

### Considerations when interpreting Red List category change

Relatively few species have experienced a genuine change in Red List category, which underestimates true numbers of changes due to: the categorical (rather than continuous) measure of extinction risk, lags in updates and biases in groups which have been reassessed. As species are assigned to Red List categories based on criteria thresholds, a species can improve or deteriorate in state (sometimes substantially) without changing Red List category and they only change category once they surpass the threshold. Even when a species does experience a genuine improvement in Red List category, a species must qualify at this level for five years before being ‘downlisted’ (IUCN, 2023). This categorisation is deliberate to facilitate assessment with lower data availability and to account for uncertainty (Mace et al., 2008), but as a result masks finer scale impacts of conservation efforts. Note it is not an issue for the Red List Index which discounts this; but is something to bear in mind generally with genuine changes in Red List category.

Birds, mammals, amphibians and warm-water reef-building corals are the only species groups that had been completely reassessed at least once, with too few corals improvements in Red List category, and justifications for improvement only available for birds and mammals. For groups that have been reassessed, updates are often infrequent; while all bird species have been reassessed every 4-5 years, other groups have been reassessed less frequently, and therefore, some recent improvements (and deteriorations) in category may not yet be reflected on the Red List or have been recorded as genuine changes. For example, the Iberian Lynx, *Lynx pardinus,* changed Red List category from Critically Endangered to Endangered in 2014, and to Vulnerable in 2023 following increases in population from 94 individuals in 2002 to 2,021 in 2023 (*Censos – Life Lynxconnect*, 2024); this was not yet recorded as a genuine category change due to delays with updating such changes for mammals.

Similarly, due to only 33 mammals experiencing improvements in Red List category, we were unable to identify actions attributed to a greater or lesser extent to these improvements, though work to review these species is currently underway. We were also unable to distinguish the primary actions that lead to improvements from all attributed actions; it would be great if assessors distinguished these primary actions as is already done for primary threats. Going forward, as more species are assessed the Green Status of Species (Akçakaya et al., 2018; IUCN, International Union for Conservation of Nature, 2021) in particular will form a key indicator to track species recovery and understand conservation impact, particularly as it compared to a pre-human baseline of impact and assesses species within spatial units within their range, so will allow a more nuanced evaluation of conservation impact.

### Conclusion

Despite inevitable data gaps and uncertainties, we demonstrate the clear exceptional value of the IUCN Red List to measure and document conservation action impact through documenting information on the actions underway for species, monitoring changes in species’ state and reporting reasons for these changes, at both global and local scales. Our results suggest that a wide range of conservation actions have been successful, particularly when targeted at specific species and locations, and have prevented extinctions of species at greatest risk. However, given relatively few species have shown signs of recovery, achieving Goal A of the Global Biodiversity Framework (CBD, 2022) will require substantially more ambitious, coordinated scaling-up of conservation interventions, particularly landscape-scale actions that benefit widely distributed species to recover global biodiversity.

## Acknowledgements

Thanks to members of the Conservation Science lab group in the Zoology Department in Cambridge and M. Hoffmann for comments on preliminary results, and to Tom Worthington for help producing a fishnet for the species richness maps. Thanks to all those who have contributed to the IUCN Red List of Threatened Species^TM^. ATS is supported through the Natural Environment Research Council’s C-CLEAR Doctoral Training Partnership (grant NE/S007164/1). For the purpose of open access, the author has applied a Creative Commons Attribution (CC BY) licence to any Author Accepted Manuscript version arising from this submission.

