## Supplementary Material for "What works to improve species conservation state? An analysis of species whose state has improved and the actions responsible"

### Supplementary Materials

Text S1. Classification of species groups.

Freshwater fish consisted of species of Actinopterygii coded with system 'freshwater'. Selected crustacea consisted species of freshwater crabs, shrimps and crayfishes and lobsters. Selected gastropods consisted species of abalones and cone snails. Selected marine fish consisted of species of Clupeiformes, blennies, boarfishes, butterflyfishes, croakers and drums, Stomiiformes, emperor breams, filefishes, groupers, grunts, Saccopharyngiformes, Carangidae, lanternfishes, Aulopiformes, louvar, Tetraodontidae, seabreams, porgies and picarels, snappers, sturgeons and paddlefishes, surgeonfishes, tangs and unicorn fishes, syngathiform fishes, seamoths, tarpons and ladyfishes, tunas, billfishes and swordfish, wrasses and parrotfishes. Warm-water reef-building corals consisted of species in the Class Hydrozoa and Family Milleporidae, or Families: Acroporidae,, Agariciidae, Astrangiidae, Astrocoeniidae, Cladocoridae, Coscinaraeidae, Diploastraeidae, Euphylliidae, Faviidae, Fungiidae, Helioporidae, Leptastreidae, Lobophylliidae, Meandrinidae, Merulinidae, Montastraeidae, Oculinidae, Oulastreidae, Plerogyridae, Plesiastreidae, Pocilloporidae, Poritidae, Psammocoridae, Rhizangiidae, Tubiporidae or Turbinoliidae; or Genus: Heterocyathus, Balanophyllia, Duncanopsammia, Heterosammia, Turbinaria, Pachyseris or Solenastrea.

Table S1. Chi-squared statistics showing for each taxonomic group and IUCN Red List category, and Red List category within each taxon, whether different groups were more or less likely to have conservation actions in place. For analysis of actions across Red List categories within taxa, this could not be evaluated for Merostomata, Myxini, Sarcopterygii and warm-water reef building corals.

| Taxon | IUCN Red List category | Residuals | p value | More/less likely |
|---|---|---|---|---|
| Aves | all | 60.56487 | <0.001*** | > |
| Amphibia | all | 50.95389 | <0.001*** | > |
| Reptilia | all | 41.35287 | <0.001*** | > |
| Warm-water reef-building corals | all | 27.55593 | <0.001*** | > |
| Merostomata | all | -0.07375 | 1 | NS |
| Sarcopterygii | all | -0.10431 | 1 | NS |
| Mammalia | all | -2.35021 | 0.600 | NS |
| Petromyzonti | all | -4.12444 | 0.001** | < |
| Myxini | all | -7.49457 | <0.001*** | < |
| Chondrichthyes | all | -10.5467 | <0.001*** | < |
| Selected marine fish | all | -16.4605 | <0.001*** | < |
| Selected gastropods | all | -22.7301 | <0.001*** | < |
| Cephalopoda | all | -26.4433 | <0.001*** | < |
| Dragonflies & damselflies | all | -28.0223 | <0.001*** | < |
| Selected crustacea | all | -42.086 | <0.001** | < |
| Freshwater fish | all | -69.1586 | <0.001*** | < |
| All | Near Threatened | 10.63879 | <0.001*** | > |
| | Vulnerable | 0.950171 | 1 | NS |
| | Endangered | -2.27432 | 0.184 | NS |
| | Critically Endangered | -10.2608 | <0.001*** | < |
| Amphibia | Near Threatened | 6.241307 | <0.001*** | > |
| | Vulnerable | 6.912726 | <0.001*** | > |
| | Endangered | -1.36612 | 1 | NS |
| | Critically Endangered | -10.3376 | <0.001*** | < |
| Aves | Near Threatened | -3.98213 | <0.001*** | < |
| | Vulnerable | 2.82727 | 0.038* | > |
| | Endangered | 0.971679 | 1 | NS |
| | Critically Endangered | 0.919876 | 1 | NS |
| Cephalopoda | Near Threatened | 1.707825 | 0.701 | NS |
| | Vulnerable | -0.68313 | 1 | NS |
| | Endangered | -0.68313 | 1 | NS |
| | Critically Endangered | -0.44096 | 1 | NS |
| Chondrichthyes | Near Threatened | -2.39078 | 0.134 | NS |
| | Vulnerable | 0.188703 | 1 | NS |

| | | | | |
|---|---|---|---|---|
| | Endangered | 0.452642 | 1 | NS |
| | Critically Endangered | 1.960376 | 0.399615 | NS |
| Dragonflies & damselflies | Near Threatened | 3.444471 | 0.005** | > |
| | Vulnerable | 1.079743 | 1 | NS |
| | Endangered | -2.28617 | 0.178 | NS |
| | Critically Endangered | -3.03665 | 0.019* | < |
| Freshwater fish | Near Threatened | 5.923018 | <0.001*** | > |
| | Vulnerable | -2.7349 | 0.050* | < |
| | Endangered | 1.777481 | 0.604 | NS |
| | Critically Endangered | -4.63233 | <0.001*** | < |
| Mammalia | Near Threatened | 0.530321 | 1 | NS |
| | Vulnerable | 1.365813 | 1 | NS |
| | Endangered | -2.34108 | 0.154 | NS |
| | Critically Endangered | 0.681128 | 1 | NS |
| Petromyzonti | Near Threatened | -1.41421 | 1 | NS |
| | Vulnerable | -1.41421 | 1 | NS |
| | Endangered | 2.165064 | 0.243 | NS |
| | Critically Endangered | 0.547723 | 1 | NS |
| Reptilia | Near Threatened | 3.939116 | <0.001*** | > |
| | Vulnerable | 2.242336 | 0.200 | NS |
| | Endangered | 0.60195 | 1 | NS |
| | Critically Endangered | -7.64468 | <0.001*** | < |
| Selected crustacea | Near Threatened | -1.21407 | 1 | NS |
| | Vulnerable | -2.62624 | 0.069 | NS |
| | Endangered | 1.080334 | 1 | NS |
| | Critically Endangered | 3.019929 | 0.020* | > |
| Selected gastropods | Near Threatened | -1.86877 | 0.493 | NS |
| | Vulnerable | -0.29053 | 1 | NS |
| | Endangered | -0.50381 | 1 | NS |
| | Critically Endangered | 3.854612 | <0.001*** | > |
| Selected marine fish | Near Threatened | 0.487054 | 1 | NS |
| | Vulnerable | -1.65371 | 0.785 | NS |
| | Endangered | 0.305829 | 1 | NS |
| | Critically Endangered | 1.613296 | 0.853 | NS |

Table S2. Chi-squared statistics showing for all taxa, each taxonomic group individually, and by IUCN Red List category whether groups are more or less likely to have each type of conservation actions in place. If actions are absent, they were not reported for any species in that group.

| Taxon | IUCN Red List category | Action type | Residuals | p value | More/less likely |
|---|---|---|---|---|---|
| All | All | In protected area | 475.64 | 0.000 | > |
| | | Legislation/Trade control | 2.25 | 0.491 | NS |
| | | Monitoring scheme | -45.33 | 0.000 | < |
| | | Species management plan | -52.71 | 0.000 | < |
| | | Awareness | -59.63 | 0.000 | < |
| | | Captive breeding | -60.33 | 0.000 | < |
| | | Area management plan | -61.53 | 0.000 | < |
| | | Harvest management plan | -62.99 | 0.000 | < |
| | | Invasive species control | -66.04 | 0.000 | < |
| | | (Re)introduced/translocated | -69.33 | 0.000 | < |
| Amphibia | All | In protected area | 232.35 | 0.000 | > |
| | | Legislation/Trade control | -19.91 | 0.000 | < |
| | | Captive breeding | -22.91 | 0.000 | < |
| | | Invasive species control | -24.94 | 0.000 | < |
| | | Awareness | -25.61 | 0.000 | < |
| | | Monitoring scheme | -26.38 | 0.000 | < |
| | | Species management plan | -26.67 | 0.000 | < |
| | | Area management plan | -28.19 | 0.000 | < |
| | | (Re)introduced/translocated | -28.40 | 0.000 | < |
| | | Harvest management plan | -29.33 | 0.000 | < |
| Aves | All | In protected area | 205.24 | 0.000 | > |
| | | Legislation/Trade control | 34.71 | 0.000 | > |
| | | Monitoring scheme | -2.86 | 0.083 | < |
| | | Species management plan | -16.76 | 0.000 | < |
| | | Awareness | -31.61 | 0.000 | < |
| | | Invasive species control | -33.68 | 0.000 | < |
| | | Captive breeding | -34.31 | 0.000 | < |
| | | (Re)introduced/translocated | -38.90 | 0.000 | < |
| | | Area management plan | -40.35 | 0.000 | < |
| | | Harvest management plan | -41.48 | 0.000 | < |
| Cephlapoda | All | In protected area | 5.94 | 0.000 | > |
| | | Legislation/Trade control | -1.82 | 0.554 | NS |
| | | Monitoring scheme | -1.82 | 0.554 | NS |
| | | Harvest management plan | -2.30 | 0.171 | NS |

| | | | | | |
|---|---|---|---|---|---|
| Chondrichthyes | All | In protected area | 36.45 | 0.000 | > |
| | | Harvest management plan | 6.15 | 0.000 | > |
| | | Legislation/Trade control | -3.64 | 0.004 | < |
| | | Monitoring scheme | -4.16 | 0.001 | < |
| | | Species management plan | -7.64 | 0.000 | < |
| | | Area management plan | -7.90 | 0.000 | < |
| | | Awareness | -8.54 | 0.000 | < |
| | | Captive breeding | -10.73 | 0.000 | < |
| Dragonflies & damselflies | All | In protected area | 128.79 | 0.000 | > |
| | | Species management plan | -17.86 | 0.000 | < |
| | | Monitoring scheme | -18.00 | 0.000 | < |
| | | Area management plan | -18.34 | 0.000 | < |
| | | Awareness | -18.34 | 0.000 | < |
| | | Legislation/Trade control | -18.61 | 0.000 | < |
| | | Invasive species control | -18.68 | 0.000 | < |
| | | Harvest management plan | -18.96 | 0.000 | < |
| Freshwater fish | All | In protected area | 156.47 | 0.000 | > |
| | | Area management plan | -4.24 | 0.000 | < |
| | | Species management plan | -15.95 | 0.000 | < |
| | | Captive breeding | -16.84 | 0.000 | < |
| | | Harvest management plan | -17.74 | 0.000 | < |
| | | Awareness | -18.94 | 0.000 | < |
| | | Monitoring scheme | -19.63 | 0.000 | < |
| | | (Re)introduced/translocated | -20.41 | 0.000 | < |
| | | Invasive species control | -20.99 | 0.000 | < |
| | | Legislation/Trade control | -21.73 | 0.000 | < |
| Mammalia | All | In protected area | 113.56 | 0.000 | > |
| | | Legislation/Trade control | 25.91 | 0.000 | > |
| | | Captive breeding | -12.21 | 0.000 | < |
| | | Awareness | -12.84 | 0.000 | < |
| | | Monitoring scheme | -15.40 | 0.000 | < |
| | | Species management plan | -17.13 | 0.000 | < |
| | | (Re)introduced/translocated | -18.91 | 0.000 | < |
| | | Harvest management plan | -19.70 | 0.000 | < |
| | | Area management plan | -20.80 | 0.000 | < |
| | | Invasive species control | -22.47 | 0.000 | < |
| Merostomata | All | Awareness | 1.29 | 1.000 | NS |
| | | (Re)introduced/translocated | -0.32 | 1.000 | NS |
| | | Harvest management plan | -0.32 | 1.000 | NS |
| | | In protected area | -0.32 | 1.000 | NS |
| | | Monitoring scheme | -0.32 | 1.000 | NS |
| Petromyzonti | All | In protected area | 1.69 | 0.910 | NS |
| | | Awareness | -0.42 | 1.000 | NS |

| | | | | | |
|---|---|---|---|---|---|
| | | Legislation/Trade control | -0.42 | 1.000 | NS |
| | | Monitoring scheme | -0.42 | 1.000 | NS |
| | | Species management plan | -0.42 | 1.000 | NS |
| Reptilia | All | In protected area | 236.57 | 0.000 | > |
| | | Legislation/Trade control | -1.85 | 1.000 | NS |
| | | Area management plan | -28.28 | 0.000 | < |
| | | Species management plan | -28.59 | 0.000 | < |
| | | Captive breeding | -28.70 | 0.000 | < |
| | | Awareness | -28.74 | 0.000 | < |
| | | Invasive species control | -29.09 | 0.000 | < |
| | | Monitoring scheme | -29.44 | 0.000 | < |
| | | (Re)introduced/translocated | -30.40 | 0.000 | < |
| | | Harvest management plan | -31.48 | 0.000 | < |
| Sarcopterygii | All | Legislation/Trade control | 1.66 | 0.586 | NS |
| | | In protected area | -0.83 | 1.000 | NS |
| | | Monitoring scheme | -0.83 | 1.000 | NS |
| Selected crustacea | All | In protected area | 45.53 | 0.000 | > |
| | | Harvest management plan | 4.87 | 2.3e-05 | > |
| | | Area management plan | -3.30 | 0.020 | < |
| | | Monitoring scheme | -4.55 | 0.000 | < |
| | | Awareness | -6.28 | 0.000 | < |
| | | Species management plan | -6.28 | 0.000 | < |
| | | (Re)introduced/translocated | -7.06 | 0.000 | < |
| | | Legislation/Trade control | -7.06 | 0.000 | < |
| | | Captive breeding | -7.85 | 0.000 | < |
| | | Invasive species control | -8.01 | 0.000 | < |
| Selected gastropods | All | In protected area | 14.49 | 0.000 | > |
| | | Harvest management plan | -1.04 | 1.000 | NS |
| | | Species management plan | -1.38 | 1.000 | NS |
| | | Monitoring scheme | -1.73 | 1.000 | NS |
| | | (Re)introduced/translocated | -2.07 | 0.615 | NS |
| | | Area management plan | -2.07 | 0.615 | NS |
| | | Captive breeding | -2.76 | 0.092 | < |
| | | Awareness | -3.45 | 0.009 | < |
| Selected marine fish | All | In protected area | 115.89 | 0.000 | > |
| | | Harvest management plan | 0.59 | 1.000 | NS |
| | | Area management plan | -9.01 | 0.000 | < |
| | | Legislation/Trade control | -12.19 | 0.000 | < |
| | | Awareness | -13.32 | 0.000 | < |
| | | Monitoring scheme | -15.16 | 0.000 | < |
| | | Species management plan | -15.73 | 0.000 | < |
| | | Captive breeding | -15.87 | 0.000 | < |

| | | | | | |
|---|---|---|---|---|---|
| | | Invasive species control | -17.56 | 0.000 | < |
| | | (Re)introduced/translocated | -17.63 | 0.000 | < |
| Warm-water reef building corals | All | In protected area | 63.91 | 0.000 | > |
| | | Legislation/Trade control | -15.90 | 0.000 | < |
| | | Captive breeding | -16.00 | 0.000 | < |
| | | Monitoring scheme | -16.00 | 0.000 | < |
| | | Species management plan | -16.00 | 0.000 | < |
| All | Near Threatened | In protected area | 12.79 | 0.000 | > |
| | | Area management plan | -0.64 | 1.000 | NS |
| | | Legislation/Trade control | -1.18 | 1.000 | NS |
| | | Harvest management plan | -1.19 | 1.000 | NS |
| | | Monitoring scheme | -2.05 | 1.000 | NS |
| | | (Re)introduced/translocated | -2.86 | 0.171 | NS |
| | | Invasive species control | -3.04 | 0.096 | NS |
| | | Species management plan | -5.31 | 4e-06 | < |
| | | Awareness | -5.75 | 0.000 | < |
| | | Captive breeding | -8.46 | 0.000 | < |
| | Vulnerable | In protected area | 2.62 | 0.350 | NS |
| | | Legislation/Trade control | 1.07 | 1.000 | NS |
| | | Harvest management plan | 0.54 | 1.000 | NS |
| | | (Re)introduced/translocated | 0.08 | 1.000 | NS |
| | | Species management plan | -0.06 | 1.000 | NS |
| | | Invasive species control | -0.64 | 1.000 | NS |
| | | Monitoring scheme | -0.72 | 1.000 | NS |
| | | Area management plan | -1.24 | 1.000 | NS |
| | | Captive breeding | -2.48 | 0.519 | NS |
| | | Awareness | -3.43 | 0.024 | < |
| | Endangered | Awareness | 1.83 | 1.000 | NS |
| | | Captive breeding | 1.52 | 1.000 | NS |
| | | Invasive species control | 0.38 | 1.000 | NS |
| | | (Re)introduced/translocated | 0.04 | 1.000 | NS |
| | | In protected area | -0.06 | 1.000 | NS |
| | | Area management plan | -0.09 | 1.000 | NS |
| | | Legislation/Trade control | -0.13 | 1.000 | NS |
| | | Species management plan | -0.61 | 1.000 | NS |
| | | Monitoring scheme | -1.46 | 1.000 | NS |
| | | Harvest management plan | -1.72 | 1.000 | NS |
| | Critically Endangered | Captive breeding | 10.42 | 0.000 | > |
| | | Awareness | 8.23 | 0.000 | > |
| | | Species management plan | 6.58 | 0.000 | > |

| | | | | | |
|---|---|---|---|---|---|
| | | Monitoring scheme | 4.82 | 5.9e-05 | > |
| | | Invasive species control | 3.62 | 0.012 | > |
| | | (Re)introduced/translocated | 2.97 | 0.120 | NS |
| | | Harvest management plan | 2.68 | 0.297 | NS |
| | | Area management plan | 2.29 | 0.878 | NS |
| | | Legislation/Trade control | 0.14 | 1.000 | NS |
| | | In protected area | -17.02 | 0.000 | < |

Table S3. Cumulative linked mixed model output, showing how (i) animal species' global population trends, (ii) genuine changes in species' Red List category and (iii) prevented decline in species' state (estimated, Green Status of Species) varies with different species' traits, threats and actions. Positive estimates indicate the variable correlates with a more favourable state, with positively and negatively correlated variables in green and orange font respectively. The below shows overall model results, models just looking at only species traits, and models for particular taxa with sufficient species and actions to evaluate independently.

| Indicator | Taxonomic Group(s) | Variable type | Variable | Estimate | SE | z value | p value |
|---|---|---|---|---|---|---|---|
| Global population trend | All comprehensively assessed taxa, excluding Least Concern Species | Traits | log(range) | -0.31 | 0.01 | -21.85 | < 0.001*** |
| | | | Red List category (linear) | -2.39 | 0.10 | -23.38 | < 0.001*** |
| | | | Red List category (quadratic) | -0.05 | 0.08 | -0.67 | 0.504 NS |
| | | | Red List category (cubic) | 0.38 | 0.07 | 5.04 | < 0.001*** |
| | | | Marine (vs terrestrial) | 0.63 | 0.46 | 1.39 | 0.163 NS |
| | | | Terrestrial and marine (vs terrestrial only) | 1.41 | 0.19 | 7.43 | < 0.001*** |

| | | | | | | |
|---|---|---|---|---|---|---|
| | Threats | Climate | 0.59 | 0.10 | 6.06 | < 0.001*** |
| | | Pollution | -0.59 | 0.10 | -5.76 | < 0.001*** |
| | | Hunting or fishing | -0.65 | 0.10 | -6.34 | < 0.001*** |
| | | Habitat loss or degradation | -1.22 | 0.09 | -13.28 | < 0.001*** |
| | Actions | Reintroduction or translocation | 1.25 | 0.18 | 7.00 | < 0.001*** |
| | | Control invasive or problematic species or disease | 0.99 | 0.14 | 7.01 | < 0.001*** |
| | | Species management plan | 0.83 | 0.13 | 6.35 | < 0.001*** |
| | | Awareness and education | 0.74 | 0.14 | 5.26 | < 0.001*** |
| | | Legislation or trade control | 0.62 | 0.11 | 5.57 | < 0.001*** |
| All comprehensively assessed taxa, including Least Concern species and only considering species traits | Traits | log(generation length) | -0.08 | 0.04 | -2.27 | 0.023 * |
| | | log(range) | -0.05 | 0.01 | -6.58 | <0.001 *** |
| | | Red List category (linear) | -2.39 | 0.11 | -21.49 | <0.001 *** |
| | | Red List category (quadratic) | 1.30 | 0.10 | 13.36 | <0.001 *** |
| | | Red List category (cubic) | -0.53 | 0.11 | -5.05 | <0.001 *** |
| | | Red List category (^4) | 0.61 | 0.09 | 6.58 | <0.001 *** |
| | | Marine (vs terrestrial) | 0.33 | 0.35 | 0.94 | 0.346 NS |
| | | Terrestrial and marine (vs terrestrial only) | 0.80 | 0.09 | 9.11 | <0.001 *** |
| Aves | Traits | log(generation length) | 1.10 | 0.17 | 6.27 | <0.001 *** |
| | | log(range) | -0.27 | 0.03 | -10.81 | <0.001 *** |
| | | Red List category (linear) | -1.08 | 0.21 | -5.10 | <0.001 *** |

| | | | | | | |
|---|---|---|---|---|---|---|
| | | Red List category (quadratic) | 0.06 | 0.18 | 0.33 | 0.744 NS |
| | | Red List category (cubic) | 0.25 | 0.17 | 1.49 | 0.136 NS |
| | Threats | Climate | 0.64 | 0.21 | 3.07 | 0.002 ** |
| | | Hunting or fishing | -0.55 | 0.19 | -2.88 | 0.004 ** |
| | | Habitat loss or degradation | -1.82 | 0.17 | -10.57 | <0.001 *** |
| | Actions | Control invasive or problematic species or disease | 1.07 | 0.21 | 5.12 | <0.001 *** |
| | | Reintroduction or translocation | 1.00 | 0.30 | 3.33 | <0.001 *** |
| | | Legislation or trade control | 0.51 | 0.18 | 2.86 | 0.004 ** |
| | | Species management plan | 0.42 | 0.20 | 2.11 | 0.035 * |
| Mammalia | Traits | log(generation length) | 0.77 | 0.21 | 3.74 | <0.001 *** |
| | | log(range) | -0.38 | 0.05 | -7.71 | <0.001 *** |
| | | Red List category (linear) | -2.07 | 0.37 | -5.62 | <0.001 *** |
| | | Red List category (quadratic) | -0.02 | 0.27 | -0.06 | 0.953 NS |
| | | Red List category (cubic) | 0.28 | 0.24 | 1.20 | 0.231 NS |
| | | Marine (vs terrestrial) | 2.59 | 0.88 | 2.96 | 0.003 ** |
| | | Terrestrial and marine (vs terrestrial only) | 1.61 | 0.70 | 2.30 | 0.022 * |
| | Threats | Problematic or invasive species or diseases | 0.58 | 0.28 | 2.07 | 0.039 * |
| | | Habitat loss or degradation | -0.99 | 0.40 | -2.46 | 0.014 * |

| | | | | | | |
|---|---|---|---|---|---|---|
| | Actions | Species management plan | 1.01 | 0.35 | 2.87 | 0.004 ** |
| | | Reintroduction or translocation | 0.97 | 0.41 | 2.36 | 0.018 * |
| | | Awareness or education | 0.91 | 0.32 | 2.81 | 0.005 ** |
| Amphibia | Traits | log(range) | -0.51 | 0.04 | -12.22 | <0.001 *** |
| | | Red List category (linear) | -4.11 | 0.26 | -15.93 | <0.001 *** |
| | | Red List category (quadratic) | -0.02 | 0.18 | -0.13 | 0.897 NS |
| | | Red List category (cubic) | 0.13 | 0.16 | 0.82 | 0.411 NS |
| | Threats | Climate | 0.76 | 0.18 | 4.17 | <0.001 *** |
| | | Pollution | -0.77 | 0.21 | -3.73 | <0.001 *** |
| | | Hunting or fishing | -0.74 | 0.35 | -2.14 | 0.032 * |
| | | Habitat loss or degradation | -1.82 | 0.28 | -6.48 | <0.001 *** |
| | Actions | Reintroduction or translocation | 2.44 | 0.61 | 4.00 | <0.001 *** |
| | | Area management plan | 2.17 | 0.69 | 3.16 | 0.002 ** |
| | | Control invasive or problematic species or disease | 1.33 | 0.39 | 3.38 | <0.001 *** |
| | | In protected area | 0.54 | 0.23 | 2.39 | 0.017 * |
| Reptilia | Traits | log(range) | -0.35 | 0.03 | -10.69 | <0.001 *** |
| | | Red List category (linear) | -2.68 | 0.25 | -10.80 | <0.001 *** |
| | | Red List category (quadratic) | 0.47 | 0.20 | 2.34 | 0.019 * |
| | | Red List category (cubic) | 0.31 | 0.20 | 1.51 | 0.130 NS |

| | | | | | | | |
|---|---|---|---|---|---|---|---|
| | | Threats | Climate | 1.33 | 0.36 | 3.71 | <0.001 *** |
| | | | Habitat loss or degradation | -1.12 | 0.21 | -5.24 | <0.001 *** |
| | | | Problematic or invasive species or diseases | -1.19 | 0.25 | -4.71 | <0.001 *** |
| | | | Pollution | -2.42 | 0.73 | -3.32 | <0.001 *** |
| | | Actions | Reintroduction or translocation | 2.76 | 0.45 | 6.11 | <0.001 *** |
| | | | Control invasive or problematic species or disease | 1.33 | 0.42 | 3.14 | 0.002 ** |
| | | | Awareness | 1.07 | 0.50 | 2.13 | 0.033 * |
| | | | Species management plan | 0.97 | 0.46 | 2.10 | 0.036 * |
| | Freshwater fish | Traits | log(range) | -0.32 | 0.04 | -8.25 | <0.001 *** |
| | | | Red List category (linear) | -2.29 | 0.24 | -9.65 | <0.001 *** |
| | | | Red List category (quadratic) | -0.37 | 0.20 | -1.90 | 0.057 NS |
| | | | Red List category (cubic) | 0.67 | 0.16 | 4.04 | <0.001 *** |
| | | Threats | Pollution | -0.43 | 0.17 | -2.57 | 0.010 * |
| | | | Habitat loss or degradation | -0.45 | 0.18 | -2.59 | 0.010 ** |
| | | | Hunting or fishing | -1.32 | 0.23 | -5.80 | <0.001 *** |
| | | Actions | Species management plan | 1.24 | 0.28 | 4.39 | <0.001 *** |
| | | | Reintroduction or translocation | 1.23 | 0.39 | 3.15 | 0.002 ** |
| Genuine change in Red List category | Amphibia, Aves and Mammalia | Traits | log(range) | 0.16 | 0.01 | 14.43 | < 0.001*** |

| | | | | | | |
|---|---|---|---|---|---|---|
| | | Red List category (linear) | 4.68 | 0.77 | 6.10 | < 0.001*** |
| | | Red List category (quadratic) | 2.12 | 0.69 | 3.06 | 0.002** |
| | | Red List category (cubic) | 1.09 | 0.48 | 2.27 | 0.023* |
| | | Red List category (^4) | 0.68 | 0.26 | 2.64 | 0.008** |
| | | Red List category (^5) | 0.13 | 0.12 | 1.09 | 0.277 NS |
| | | Marine (vs terrestrial) | 1.56 | 0.49 | 3.21 | 0.001** |
| | | Terrestrial and marine (vs terrestrial only) | 0.33 | 0.15 | 2.16 | 0.031* |
| | Threats | Climate | -0.97 | 0.08 | -12.61 | < 0.001*** |
| | | Hunting or fishing | -0.73 | 0.08 | -8.87 | < 0.001*** |
| | | Invasive or problematic species or disease | -1.11 | 0.07 | -15.88 | < 0.001*** |
| | | Habitat loss or degradation | -1.45 | 0.09 | -16.56 | < 0.001*** |
| | Actions | Reintroduction or translocation | 2.53 | 0.32 | 7.91 | < 0.001*** |
| | | Control invasive or problematic species or disease | 0.45 | 0.18 | 2.48 | 0.013* |
| | | Legislation or trade control | -0.17 | 0.09 | -1.98 | 0.048* |
| | | Awareness and education | -0.31 | 0.14 | -2.21 | 0.027* |
| | | Species management plan | -0.59 | 0.12 | -4.96 | < 0.001*** |
| Amphibia, Aves, Mammalia, only considering species traits | Traits | log(generation length) | -0.41 | 0.07 | -5.63 | <0.001 *** |
| | | log(range) | 0.03 | 0.02 | 2.19 | 0.028 * |
| | | Red List category (linear) | 5.29 | 0.71 | 7.44 | <0.001 *** |

| | | | | | | |
|---|---|---|---|---|---|---|
| | | Red List category (quadratic) | 3.68 | 0.64 | 5.73 | <0.001 *** |
| | | Red List category (cubic) | 0.37 | 0.45 | 0.83 | 0.406 NS |
| | | Red List category (^4) | 0.10 | 0.26 | 0.38 | 0.701 NS |
| | | Red List category (^5) | -0.43 | 0.15 | -2.84 | 0.004 ** |
| | | Marine (vs terrestrial) | 2.28 | 0.56 | 4.11 | <0.001 *** |
| | | Terrestrial and marine (vs terrestrial only) | -0.15 | 0.16 | -0.96 | 0.335 NS |
| Amphibia | Traits | log(range) | 0.28 | 0.02 | 16.73 | <0.001 *** |
| | | Red List category (linear) | 2.16 | 0.14 | 15.24 | <0.001 *** |
| | | Red List category (quadratic) | 0.20 | 0.11 | 1.74 | 0.082 NS |
| | | Red List category (cubic) | -0.42 | 0.12 | -3.49 | <0.001 *** |
| | | Red List category (^4) | 0.17 | 0.12 | 1.42 | 0.155 NS |
| | Threats | Hunting or fishing | -0.50 | 0.14 | -3.57 | <0.001 *** |
| | | Habitat loss or degradation | -1.00 | 0.13 | -7.69 | <0.001 *** |
| | | Climate | -1.21 | 0.10 | -12.03 | <0.001 *** |
| | | Problematic or invasive species or diseases | -1.63 | 0.09 | -17.38 | <0.001 *** |
| | Actions | In protected area | 0.40 | 0.11 | 3.59 | <0.001 *** |
| | | Species management plan | -1.45 | 0.29 | -5.00 | <0.001 *** |
| Aves | Traits | Red List category (linear) | 3.43 | 0.23 | 14.90 | <0.001 *** |
| | | Red List category (quadratic) | 1.23 | 0.20 | 6.07 | <0.001 *** |

| | | | | | | |
|---|---|---|---|---|---|---|
| | | Red List category (cubic) | 1.29 | 0.18 | 7.17 | <0.001 *** |
| | | Red List category (^4) | 0.33 | 0.16 | 2.12 | 0.034 * |
| | Threats | Problematic or invasive species or diseases | -0.51 | 0.14 | -3.53 | <0.001 *** |
| | | Hunting or fishing | -0.86 | 0.13 | -6.84 | <0.001 *** |
| | | Habitat loss or degradation | -2.87 | 0.13 | -21.81 | <0.001 *** |
| | Actions | Reintroduction or translocation | 2.66 | 0.47 | 5.71 | <0.001 *** |
| | | Legislation or trade control | 0.31 | 0.11 | 2.70 | 0.007 ** |
| | | Control invasive or problematic species or disease | -0.50 | 0.24 | -2.09 | 0.036 * |
| | | Species management plan | -0.53 | 0.15 | -3.59 | 0.000 *** |
| | | Awareness or education | -0.57 | 0.20 | -2.90 | 0.004 ** |
| Mammalia | Traits | log(range) | 0.10 | 0.04 | 2.72 | 0.007 ** |
| | | Red List category (linear) | 2.21 | 0.35 | 6.31 | <0.001 *** |
| | | Red List category (quadratic) | 1.28 | 0.28 | 4.55 | <0.001 *** |
| | | Red List category (cubic) | -0.25 | 0.23 | -1.13 | 0.261 NS |
| | | Red List category (^4) | 0.40 | 0.20 | 2.03 | 0.042 * |
| | | Marine (vs terrestrial) | 2.36 | 0.64 | 3.69 | <0.001 *** |
| | | Terrestrial and marine (vs terrestrial only) | -1.28 | 0.41 | -3.11 | 0.002 ** |
| | Threats | Habitat loss or degradation | -0.98 | 0.24 | -4.12 | <0.001 *** |
| | | Hunting or fishing | -1.29 | 0.20 | -6.56 | <0.001 *** |

| | | | | | | | |
|---|---|---|---|---|---|---|---|
| | | Actions | Reintroduction or translocation | 3.43 | 0.53 | 6.41 | <0.001 *** |
| | | | In protected area | -0.60 | 0.18 | -3.37 | <0.001 *** |
| | | | Area management plan | -0.86 | 0.36 | -2.40 | 0.016 * |
| | | | Awareness or education | -0.89 | 0.25 | -3.51 | <0.001 *** |
| | | | Control invasive or problematic species or disease | -1.04 | 0.50 | -2.07 | 0.039 * |
| Prevented declines in state in spatial units (Green Status of Species) | All animals with published Green Status assessments | | (Intercept) | -3.62 | 0.65 | -5.61 | < 0.001*** |
| | | Traits | Counterfactual state (linear) | -1.12 | 0.55 | -2.04 | 0.041* |
| | | | Counterfactual state (quadratic) | 0.70 | 0.39 | 1.80 | 0.072 NS |
| | | Actions | Species management plan | 2.57 | 0.68 | 3.79 | < 0.001*** |
| | | | Reintroduction or translocation | 2.51 | 0.87 | 2.88 | 0.004* |
| | | | Area management plan | 1.86 | 0.45 | 4.13 | < 0.001*** |
| | | | Control invasive or problematic species or disease | 1.72 | 0.88 | 1.96 | 0.050* |
| | | | In protected area | 1.46 | 0.53 | 2.75 | 0.006** |

Table S4. Chi-squared post-hoc tests of actions which led to improvements in species' Red List category for birds and mammals, and birds only, compared to the other actions the same set of species.

| **Action** | **Birds and mammals** | | | **Birds** | | |
|---|---|---|---|---|---|---|
| | Residuals | p value | More/less likely | Residuals | p value | More/less likely |
| Reintroduced or translocated | 4.24 | 0.000 | >*** | 4.00477 | 0.001242 | >** |
| Area management plan | 2.89 | 0.078 | NS | 4.00477 | 0.001242 | >** |
| In protected area | 2.89 | 0.078 | NS | 1.649023 | 1 | NS |
| Control of invasive or problematic species or diseases | 2.35 | 0.377 | NS | 3.121365 | 0.036003 | > |
| Species management plan | 0.73 | 1.000 | NS | 0.471149 | 1 | NS |
| Captive breeding | -1.43 | 1.000 | NS | -1.29566 | 1 | NS |
| Awareness or education | -2.24 | 0.501 | NS | -2.17907 | 0.586535 | NS |
| Harvest management plan | -2.51 | 0.241 | NS | -2.768 | 0.112802 | NS |
| Legislation or trade control | -2.51 | 0.241 | NS | -3.06247 | 0.043904 | <* |
| Monitoring scheme | -4.40 | 0.000 | <*** | -3.94588 | 0.00159 | <** |

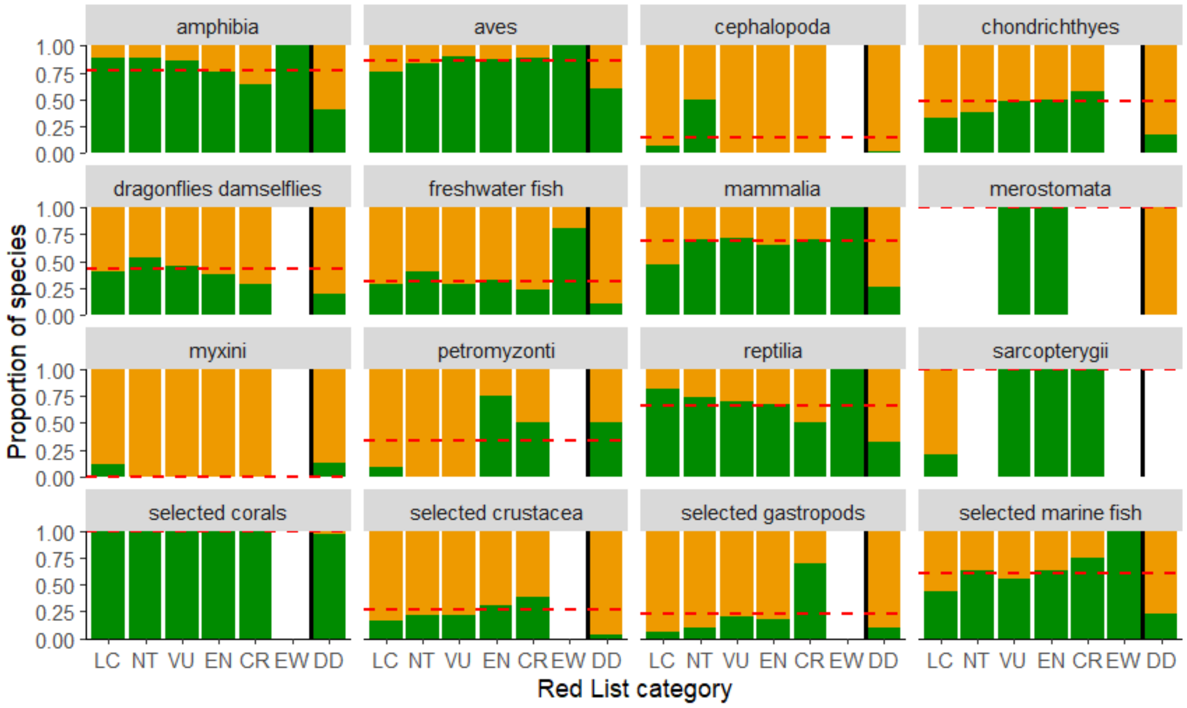


Figure S1. The proportion of species with reported conservation actions in place across different animal taxonomic groups that have been comprehensively assessed on the IUCN Red List, split by their Red List category. The dashed red line indicates the mean proportion across all species.

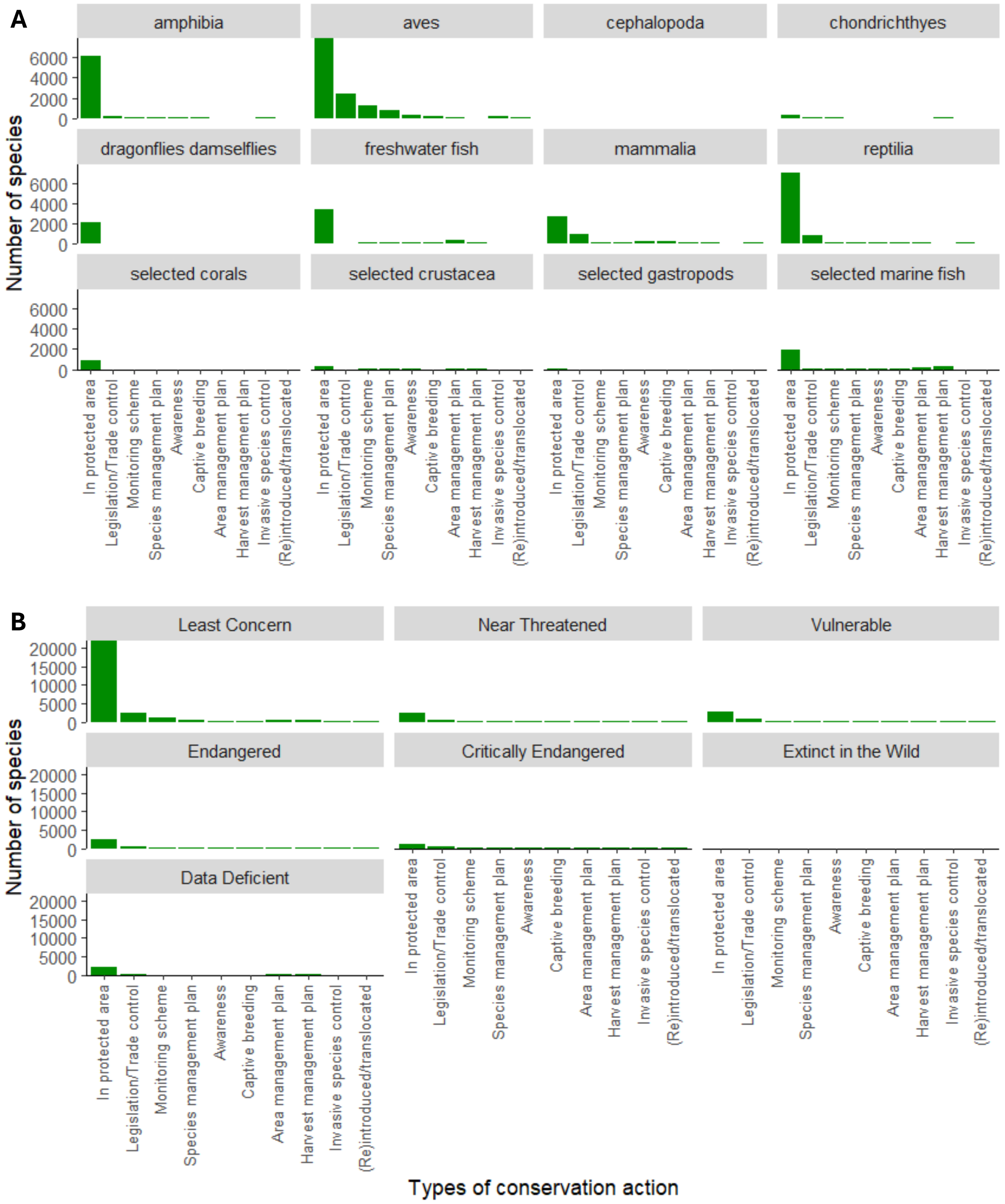


Figure S2. The percentage of species in animal groups which have been comprehensively assessed by the IUCN Red List that have each type of conservation action in place, (A) split by taxa that have been comprehensively assessed in the IUCN Red List (more than 80% assessed, excluding Cephalopoda, Merostomata, Myxini, Petromyzontid and Sarcopterygii) and (B) split by IUCN Red List category.
